# Cooperative Allostery and Structural Dynamics of Streptavidin at Cryogenic- and Ambient-temperature

**DOI:** 10.1101/2021.09.23.461567

**Authors:** Esra Ayan, Busra Yuksel, Ebru Destan, Fatma Betul Ertem, Gunseli Yildirim, Meryem Eren, Oleksandr M. Yefanov, Anton Barty, Alexandra Tolstikova, Gihan K. Ketawala, Sabine Botha, E. Han Dao, Brandon Hayes, Mengning Liang, Matthew H. Seaberg, Mark S. Hunter, Alex Batyuk, Valerio Mariani, Zhen Su, Frederic Poitevin, Chun Hong Yoon, Christopher Kupitz, Aina Cohen, Tzanko Doukov, Raymond G. Sierra, Çağdaş Dağ, Hasan DeMirci

## Abstract

Multimeric protein assemblies are abundant in nature. Streptavidin is an attractive protein that provides a paradigm system to investigate the intra- and intermolecular interactions of multimeric protein complexes. Also, it offers a versatile tool for biotechnological applications. Here, we present two apo-streptavidin structures, the first one is an ambient temperature Serial Femtosecond X-ray crystal (Apo-SFX) structure at 1.7 Å resolution and the second one is a cryogenic crystal structure (Apo-Cryo) at 1.1 Å resolution. These structures are mostly in agreement with previous structural data. Combined with computational analysis, these structures provide invaluable information about structural dynamics of apo streptavidin. Collectively, these data further reveal a novel cooperative allostery of streptavidin which binds to substrate via water molecules that provide a polar interaction network and mimics the substrate biotin which displays one of the strongest affinities found in nature.

## INTRODUCTION

Core streptavidin is a biotin-binding protein and has a homotetrameric structure consisting of 126 amino acid residues in each identical monomer [1]. The secondary structure of streptavidin involves the formation of an anti-parallel eight-stranded β-barrel tertiary structure [2]. Trp120 residue in each monomer is responsible for the linkage of the monomer to its neighboring subunits. Tetramers can be seen as composed of two functional dimers [3]. Although the function of streptavidin is not well-known, previous studies suggest that streptavidin may play a role in microbial defense mechanisms [4].

For many years, the strong non-covalent bond between biotin and streptavidin has attracted the attention of scientists in basic and applied sciences. Theoretically, the strength of streptavidin-biotin conjugation is known by its pinpoint specification and femtomolar affinity (Kd = 10^-13 to 10^-15 M) [5]. This affinity is an order of magnitude higher than prominent antigen-antibody interactions [6]. This specific affinity of the streptavidin-biotin complex contains remarkable properties. First, there is a high shape-complementarity between the binding pocket and biotin, and this affinity is strengthened by hydrogen bonds with specific residues, i.e. Asn23, Ser42, Tyr43, Ser45, Asn49, Ser88, Thr90, and Asp128 [7]. Additionally, in the hydrophobic binding pocket, numerous Van der Waals contacts with biotin make this one of the strongest non-covalent interactions with molecular dynamics studies highlighting the non-polar Van der Waals contribution of tryptophan residues e.g. Trp79, Trp92, Trp108, and Trp120 rather than electrostatic forces [8,9,3]. Protein regions which are involved in protein-protein and protein-target interactions have high plasticity and are often highly flexible. In addition, these regions are usually located on the solvent exposed surface of the protein. Apo-state streptavidin loop (L3/4; residues 45–52) **(Supplementary Fig. 1)** is the crucial segment of streptavidin that interacts with biotin and regulates binding action [10].

Previous studies have contributed significantly to our discovery-based knowledge about the streptavidin-biotin structure **(Supplementary Fig. 2)** and provided innovations in biotechnology, including a drug delivery method and catalysis, detection and labeling of molecules [11,12,13,14,15]. Furthermore, streptavidin provided numerous pragmatic platforms as a tool for orienting and delivering signals from proteins [16]. New applications are added continually to develop new organic nano-molecules. The structural dynamics of streptavidin and its interactions with small molecules demonstrate that we have a greater understanding of its structure-function relationship, which makes it an easy to “plug and play” type of molecule [15]. Streptavidin is a molecule characterized by its interactions with a wide range of biotinylated molecules and is widely used in biotechnology. Its binding with biotinylated molecules is diffusion-limited and has a remarkably high binding efficiency. Streptavidin can form new nano-assemblies by using biotinylated non-biological building blocks as well as organic molecules such as sugar, protein, and nucleic acids [15]. Tetrameric streptavidin displays non-symmetrical subunit structures in the L3/4 region. The mutated residues (N23A/S27D/S45A) favor the open conformation of the L3/4 and further decrease the biotin-binding affinity [17]. Engineered divalent cis- and trans-plane-dependent streptavidin demonstrated that the overlay of wild-type and the mutated residues may also cause a decrease in the number of polar linkages interacting with biotin [14]. These mutated residues offer a potential target for further modulating streptavidin properties.

There are conflicting hypotheses on the cooperativity of the streptavidin-biotin interaction. Some suggest that these interactions are cooperative [18, 19] while others suggest noncooperative binding based on the holo- and apo-streptavidin-biotin complexes [20]. Along with two opposing views, it was also suggested that the streptavidin-biotin interaction showed positive cooperativity, and the cooperativity of the tetramer was accompanied by closeness of the streptavidin upon biotin-binding [21, 22]. Later on, this phenomenon was named “cooperative allosterism” [23]. Streptavidin-biotin complex has numerous applications and many of them are investigated structurally. There are existing cryogenic apo-state structures in the literature [24]; however, temperature artifacts could introduce errors, preventing successful computational predictions [25]. In addition, cryogenic temperature may perturb the overall protein backbone fold [26]. Therefore, studies have recommended greater caution when referring only to cryogenic structures [24, 25]. In order to reveal the underpinnings of this elegant system, it is essential to examine the details of structural dynamics of the apo-state at near-physiological temperatures which are currently a missing reference point.

Streptavidin structure at ambient temperature will provide further information to explore the structural dynamics of this protein in more detail **(Supplementary Fig. 2)**. We obtained the radiation-damage-free ambient temperature Apo-SFX structure of streptavidin at 1.7 Å resolution during the first ever remote data collection at the Macromolecular Femtosecond Crystallography (MFX) instrument at Linac Coherent Light Source (LCLS) [27]. The diffraction data are collected by using the new generation ePix10k2M detector which has a much improved dynamic range and lower noise than the previous family of detectors [28] **(Supplementary Fig. 3)**. In addition to our Apo-SFX structure, we also present Apo-Cryo structure at 1.1 Å resolution for structural comparison. We re-evaluated the tetrameric structure of streptavidin by Gaussian Network Model (GNM) analysis of the protein’s dynamics [29] by inspecting the Apo-SFX and Holo-SFX (PDB ID: 5JD2) structures. The new Apo-SFX structure, which has higher resolution and improved electron density quality compared to previous ambient temperature apo-structures, and Holo-SFX structure were determined from data collected at the same X-ray source. Thus, temperature- and instrument-related differences and variables are minimized within these structures for accurate comparison in GNM analysis. To highlight the structural dynamics of streptavidin, we investigated the polar interaction network, number of coordinated water molecules, thermal ellipsoid structures and electrostatic surface models for both ambient and cryogenic structures. Our data presented here provide a novel cooperative allosteric model for streptavidin biotin interactions.

## RESULTS

### Ambient-temperature X-Ray Free-Electron Laser (XFEL) and cryogenic synchrotron structures of Streptavidin revealed alternate conformations of the binding pocket

We determined the first radiation-damage-free SFX crystal structure of apo-streptavidin at 1.7 Å resolution **(Fig. 1a,b & Table 1)**. The diffraction data were collected during the first remote beamtime at the MFX instrument of the LCLS at SLAC National Laboratory, Menlo Park, CA [27]. The second crystal structure of apo-streptavidin was determined at a 1.1 Å resolution at cryogenic temperatures at beamline 12-2 of the Stanford Synchrotron Radiation Lightsource (SSRL) in Menlo Park, CA **(Supplementary Fig. 4a-c & Table 1)**. Subunits of the two tetrameric crystal structures at ambient and cryogenic temperatures were superposed with an overall RMSD of 0.17 Å and 0.13 Å, respectively **(Fig. 1c & Supplementary Fig. 4c)**. Conformational changes were observed around the loop regions of these monomers.

**Fig. 1:**
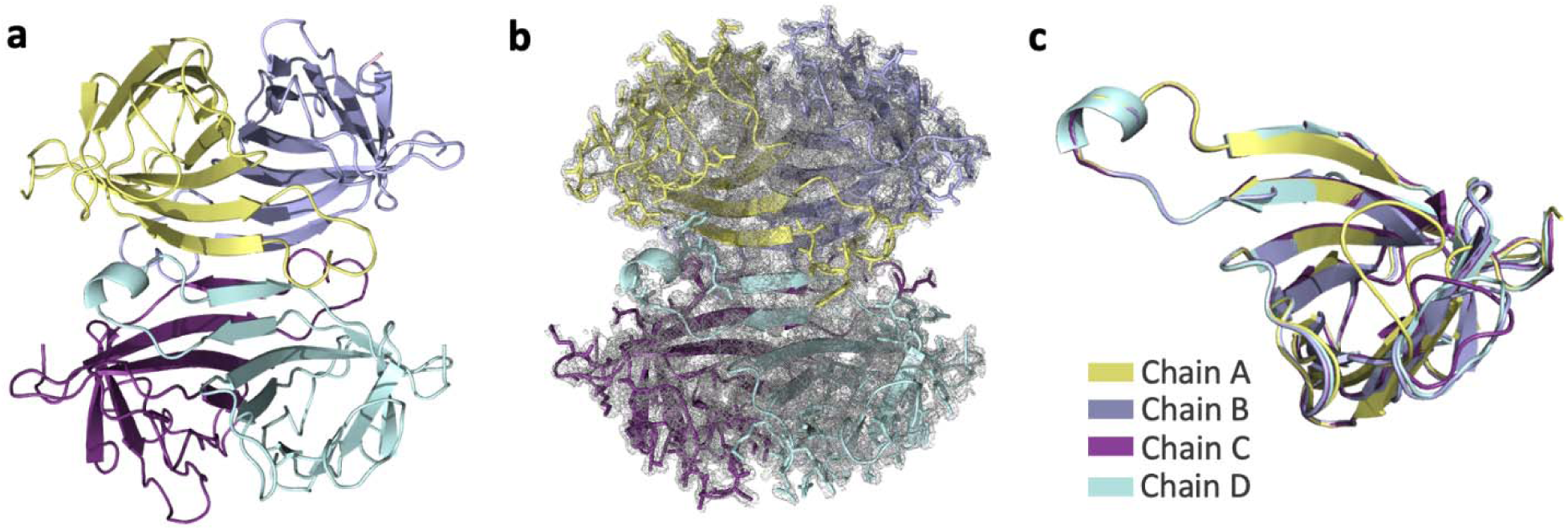
Apo-SFX structure of streptavidin. **(a)** The Apo structure of streptavidin is colored based on each chain. **(b)** 2*F*o-*F*c simulated annealing-omit map at 1 sigma level is colored in gray. **(c)** Each chain of streptavidin is superposed with an overall RMSD of 0.177 Å.

**Table 1:**
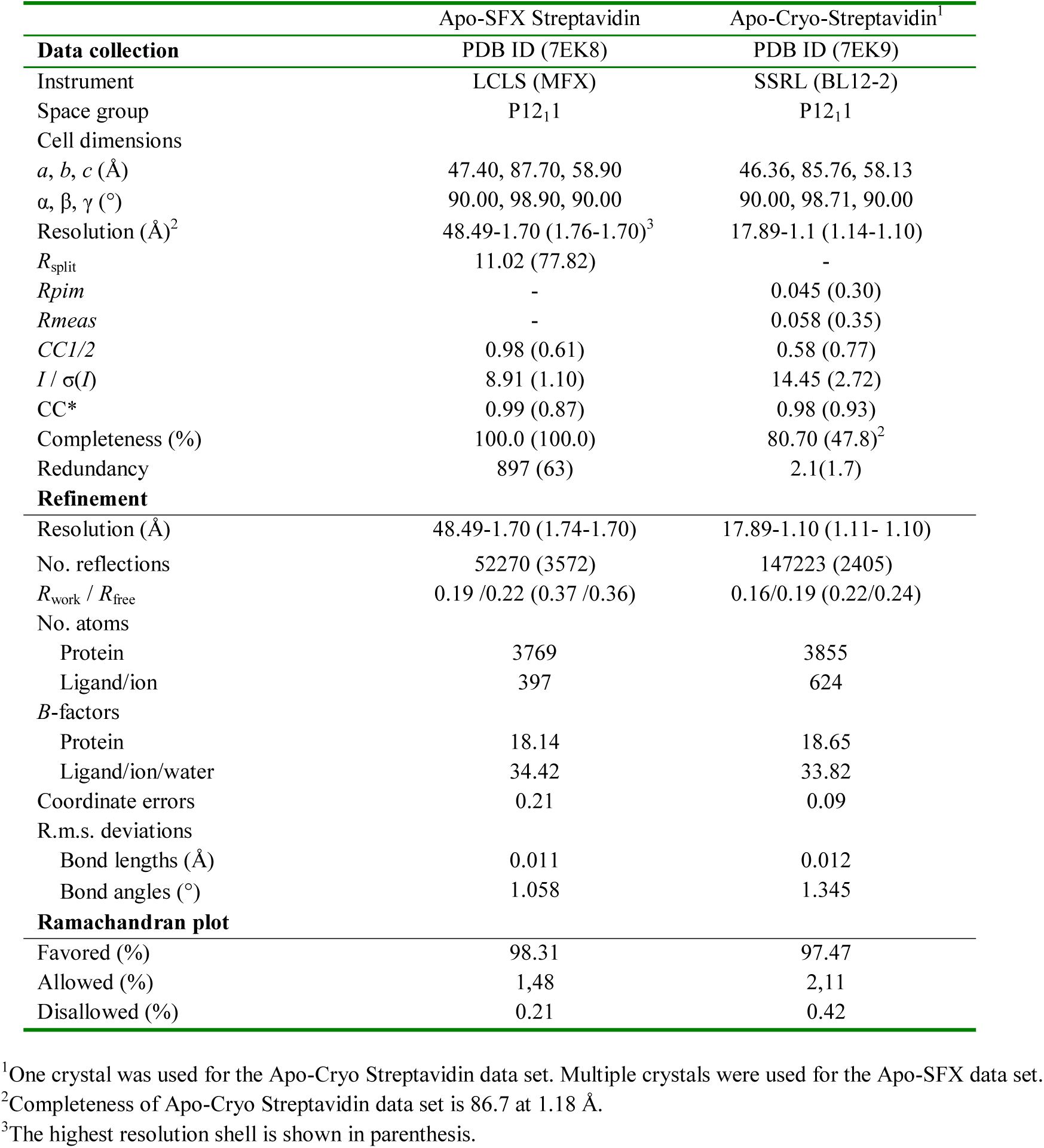
Data collection and refinement statistics.

|  | Apo-SFX Streptavidin | Apo-Cryo-Streptavidin<sup>1</sup> |
| --- | --- | --- |
| **Data collection** | PDB ID (7EK8) | PDB ID (7EK9) |
| Instrument | LCLS (MFX) | SSRL (BL12-2) |
| Space group | P12<sub>1</sub>1 | P12<sub>1</sub>1 |
| Cell dimensions |  |  |
| *a*, *b*, *c* (Å) | 47.40, 87.70, 58.90 | 46.36, 85.76, 58.13 |
| $\alpha$, $\beta$, $\gamma$ (°) | 90.00, 98.90, 90.00 | 90.00, 98.71, 90.00 |
| Resolution (Å)<sup>2</sup> | 48.49-1.70 (1.76-1.70)<sup>3</sup> | 17.89-1.1 (1.14-1.10) |
| *R*<sub>split</sub> | 11.02 (77.82) | - |
| *R*<sub>pim</sub> | - | 0.045 (0.30) |
| *R*<sub>meas</sub> | - | 0.058 (0.35) |
| *CC*1/2 | 0.98 (0.61) | 0.58 (0.77) |
| *I* / $\sigma$(*I*) | 8.91 (1.10) | 14.45 (2.72) |
| *CC* * | 0.99 (0.87) | 0.98 (0.93) |
| Completeness (%) | 100.0 (100.0) | 80.70 (47.8)<sup>2</sup> |
| Redundancy | 897 (63) | 2.1(1.7) |
| **Refinement** |  |  |
| Resolution (Å) | 48.49-1.70 (1.74-1.70) | 17.89-1.10 (1.11- 1.10) |
| No. reflections | 52270 (3572) | 147223 (2405) |
| *R*<sub>work</sub> / *R*<sub>free</sub> | 0.19 /0.22 (0.37 /0.36) | 0.16/0.19 (0.22/0.24) |
| No. atoms |  |  |
| Protein | 3769 | 3855 |
| Ligand/ion | 397 | 624 |
| *B*-factors |  |  |
| Protein | 18.14 | 18.65 |
| Ligand/ion/water | 34.42 | 33.82 |
| Coordinate errors | 0.21 | 0.09 |
| R.m.s. deviations |  |  |
| Bond lengths (Å) | 0.011 | 0.012 |
| Bond angles (°) | 1.058 | 1.345 |
| **Ramachandran plot** |  |  |
| Favored (%) | 98.31 | 97.47 |
| Allowed (%) | 1.48 | 2.11 |
| Disallowed (%) | 0.21 | 0.42 |

<sup>1</sup>One crystal was used for the Apo-Cryo Streptavidin data set. Multiple crystals were used for the Apo-SFX data set.
<sup>2</sup>Completeness of Apo-Cryo Streptavidin data set is 86.7 at 1.18 Å.
<sup>3</sup>The highest resolution shell is shown in parenthesis.

Superposition of our Apo-SFX and Apo-Cryo structures revealed very similar conformations of the substrate-binding sites with an overall RMSD of 0.30 Å **(Fig. 2a-d & Supplementary Table 1)**. We observed additional electron density that belongs to the residues in the ligand binding site region which were not modeled in previous studies (**Fig. 1a-c & Supplementary Fig. 4a-c & Supplementary Fig. 5, Supplementary Fig. 6**). First we compared the ambient temperature synchrotron apo-structure of streptavidin (1SWB) with our Apo-SFX structure [30]. 1SWB was determined at 1.85 Å resolution with an R_work_ of 0.17 and R_free_ of 0.25 while the Apo-SFX structure was determined at 1.7 Å resolution with an R_work_ of 0.19 and R_free_ of 0.22. The 1SWB structure has missing residues in chain B at positions 45-48, and in chain C and D at positions 46-48, as mentioned in **Supplementary Fig. 5**. Moreover, 1SWB was observed with lack of electron density at residues Gln24, Lue25, Val47, Glu51, and Arg53 in different chains, thus this structure was not suitable for applying GNM analysis. Moreover for a fair comparison, we compared the electron density of binding site residues between our Apo-SFX structure and the latest synchrotron cryo-structure of apo-streptavidin (PDB ID: 3RY1) [31], and found that the electron density of the loops, unfortunately, was not enhanced significantly **(Supplementary Fig. 6)**. Overall, 3RY1 was determined at 1.03 Å resolution, while Apo-SFX was determined at a lower resolution. On the other hand, in our Apo-SFX data, some of the binding site residues’ electron density enhanced, and continuous electron density without alternate side chains conformations was observed. In particular, chain A within the Apo-SFX structure between Ile30-Thr40, Ala46-Arg53, and Glu44 have clearer electron density and precise side-chain conformations compared to the 3RY1 structure as indicated in the following figure. In chain B, between Asn23-Gly26, Phe29-Leu39, Ser45-Gly48, Glu51-Val55, and Thr42 there is more precise electron density and a lack of alternate side-chain conformations for the Apo-SFX structure. In chain C Asn23-Leu25, Ile30, The32, Ala35, Glu44, and Glu51 have clearer and continuous electron density with less alternate conformations within the Apo-SFX structure, however between Ala46-Ala50 and Arg53, the electron density is clearer at 3RY1, but has similar conformations to Apo-SFX. Similarly, in chain D the Apo-SFX structure has a clearer density and more precise conformations for Asn23-Leu25, Ile30, The32, Ala35-Ala38 (but not Asp36) and Thr42, however, for Glu44-Ser52 the 3RY1 structure has clearer electron density but similar conformations to Apo-SFX.

**Fig. 2:**
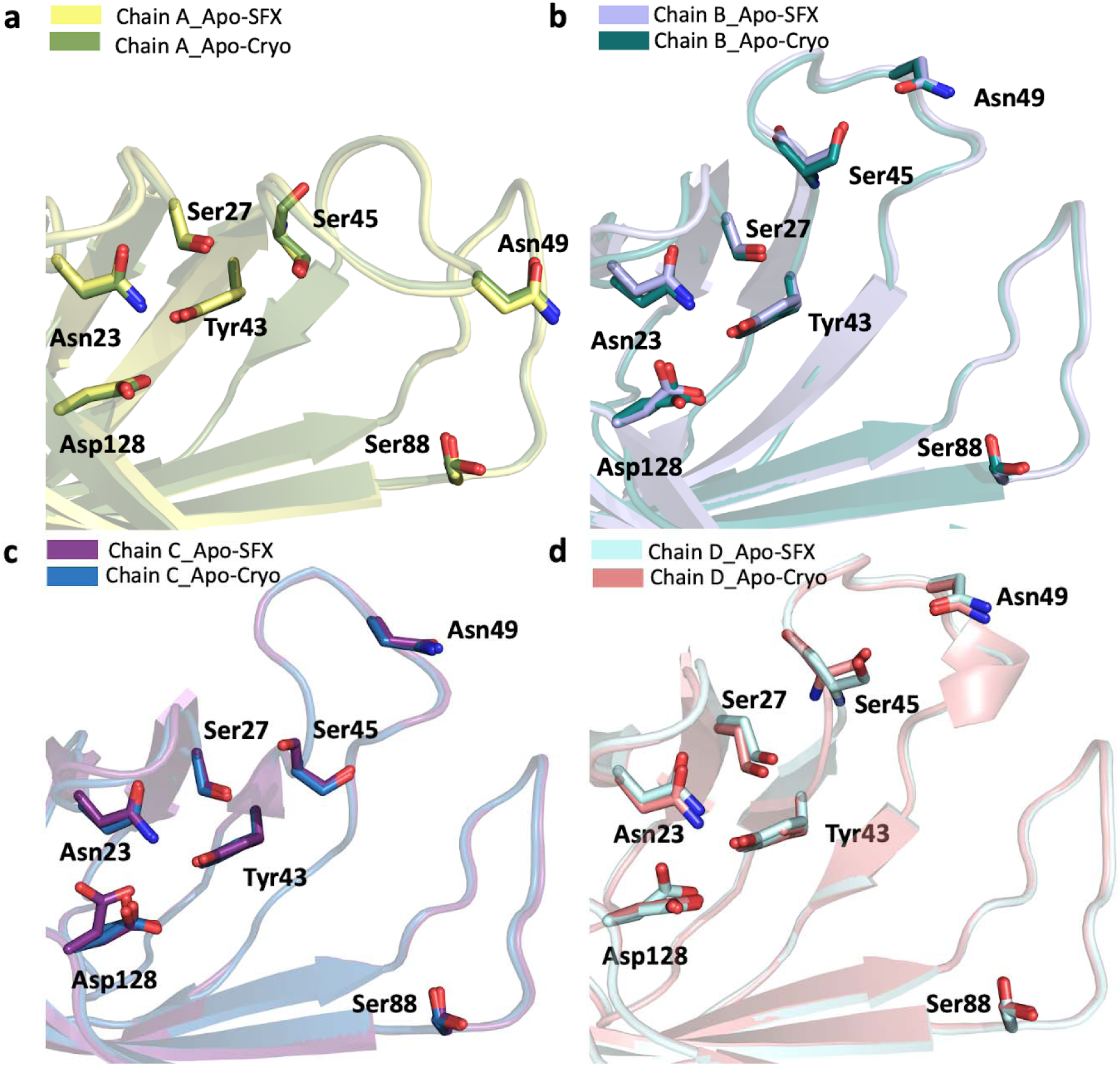
Biotin binding site comparison of Apo-SFX and Apo-Cryo structures of streptavidin. Chain A-D of Apo-SFX is superposed with Apo-Cryo structure in panel a-d, respectively (Supp Table 1). There are no significant conformational differences between the two structures.

The non-covalent interactions between biotin and streptavidin in the binding pocket residues (Asn23, Ser27, Tyr43, Ser45, Asn49, Ser88, Asp128) play an essential role in making the structure more stable [7]. To explore the noncovalent interactions in the binding pocket, hydrogen bonds were mapped between the active site and neighboring residues **(Supplementary Fig. 7a,b)**. In doing so, the superposition of Apo-SFX and Apo-CryoEM (PDB_ID:6J6K) structures revealed the differences between the substrate-binding pocket residues **(Supplementary Fig. 8a-d & Supplementary Table 1)**. Especially the residues Ser45 and Asn49 around the L3/4 region showed significant conformational differences. These residues are important for the noncovalent interactions with a biotin substrate. Interestingly, this conformational change is more prominent for chains B, C, and D. Conformational differences of L3/4 indicate the non-symmetric binding of biotin with monomers **(Supplementary Fig 9a-d)**. The superposition of our Apo-SFX structure was performed by using the Holo-CryoEM structure **(PDB ID: 6J6J) (Supplementary Fig. 9a-d & Supplementary Table 1)** and Holo-SFX structure **(Supplementary Fig. 10a-d, Fig 3a-d, Supplementary Table 2).** Additionally, we superposed Apo-Cryo structure with the Holo-SFX structure **(Supplementary Fig. 11a-d, Supplementary Table 2)**. As expected, conformational changes were found in the binding pocket. Furthermore, we observed the minor conformational changes which are less than 1 Å at the residues Ala65, Thr66, Asp77, Ala100, Glu101, Glu116, Ala117, Asp128, Lys132, Val133 and Lys134 correlated with previous mutation studies to elucidate its dynamic application [19,32,33,34,35] **(Supplementary Fig. 12a-d)**.

**Fig. 3:**
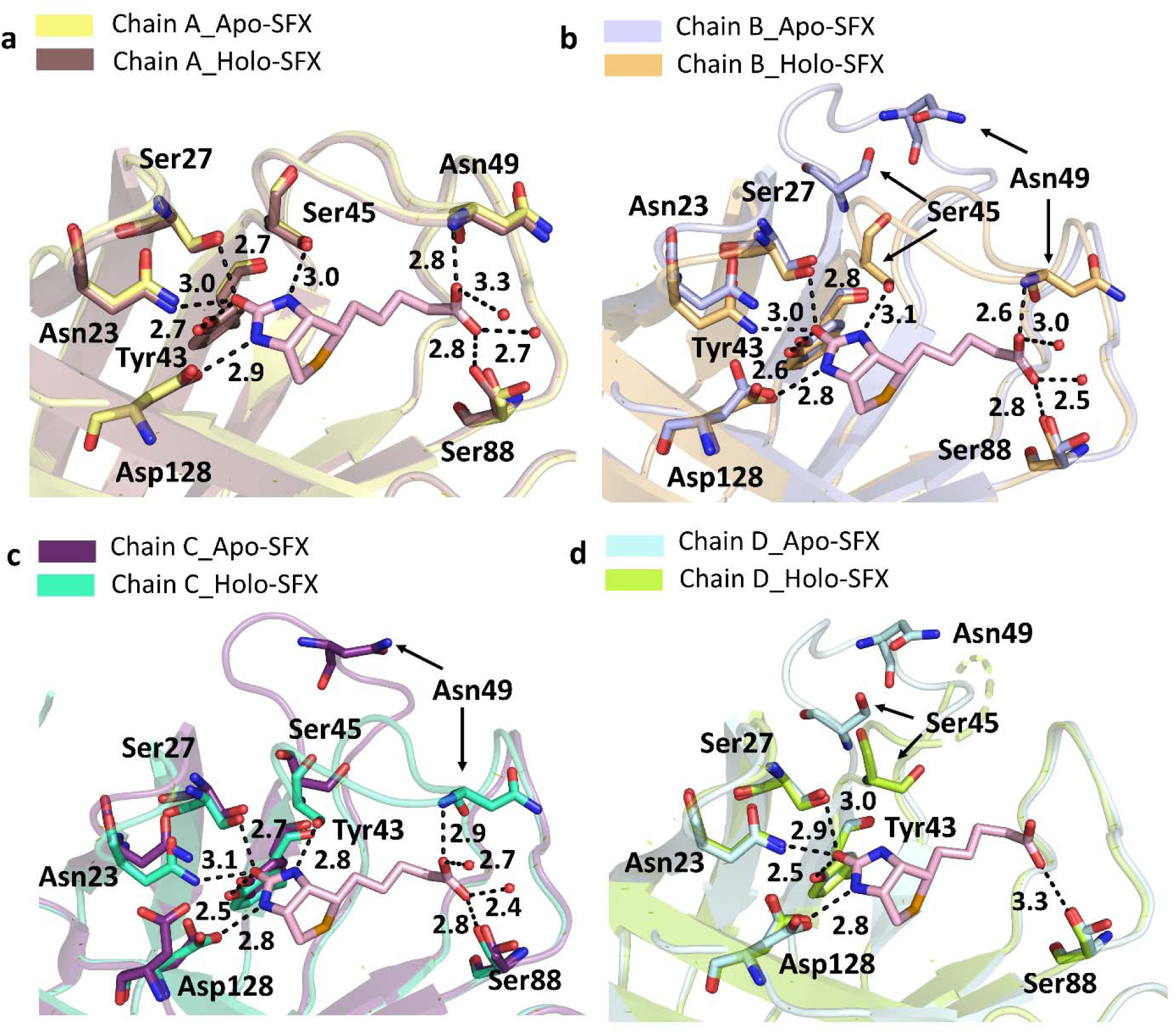
Superposition of the biotin-binding sites for each chain of the Apo-SFX and Holo-SFX structures. Chain A-D of Apo-SFX is superposed with the Holo-SFX (PDB ID:5JD2) structure of streptavidin in panel a-d, respectively (Supp Table 1). The L3/4 opening as a “lid” without selenobiotin binding. Binding of selenobiotin is not symmetric for all four monomers which represent cooperativity. Selenobiotin and water molecules were represented by light pink sticks and red-colored spheres, respectively. Hydrogen bonds are shown with black dashed lines and their corresponding distance as a unit of Angstrom (Å).

To provide more detailed information about the binding pocket conformation and interactions of streptavidin in all three structures **(Supplementary Fig. 13a-c)**, water molecules around the binding pocket of each chain were investigated **(Fig. 4a-c)**. Apo structures have a wider opening for ligand-binding at B, C, and D chains compared to the Holo-SFX structure. On the other hand, chain A displayed disrupted symmetry and an alternate conformation with a closed state similar to the Holo-SFX conformation **(Fig. 4a-c)**. Moreover, our data reveal that chain A, which has the highest number of coordinated water molecules, favors the closed state of the flexible loop due to polar interactions. Similarly, the open-loop conformation correlates with a gradually reduced number of coordinated water molecules and polar contacts in the binding pocket **(Fig. 5a-d)**. Our data suggest that coordinated water molecules in the binding pocket and polar interaction network have a key role in cooperativity.

**Fig. 4:**
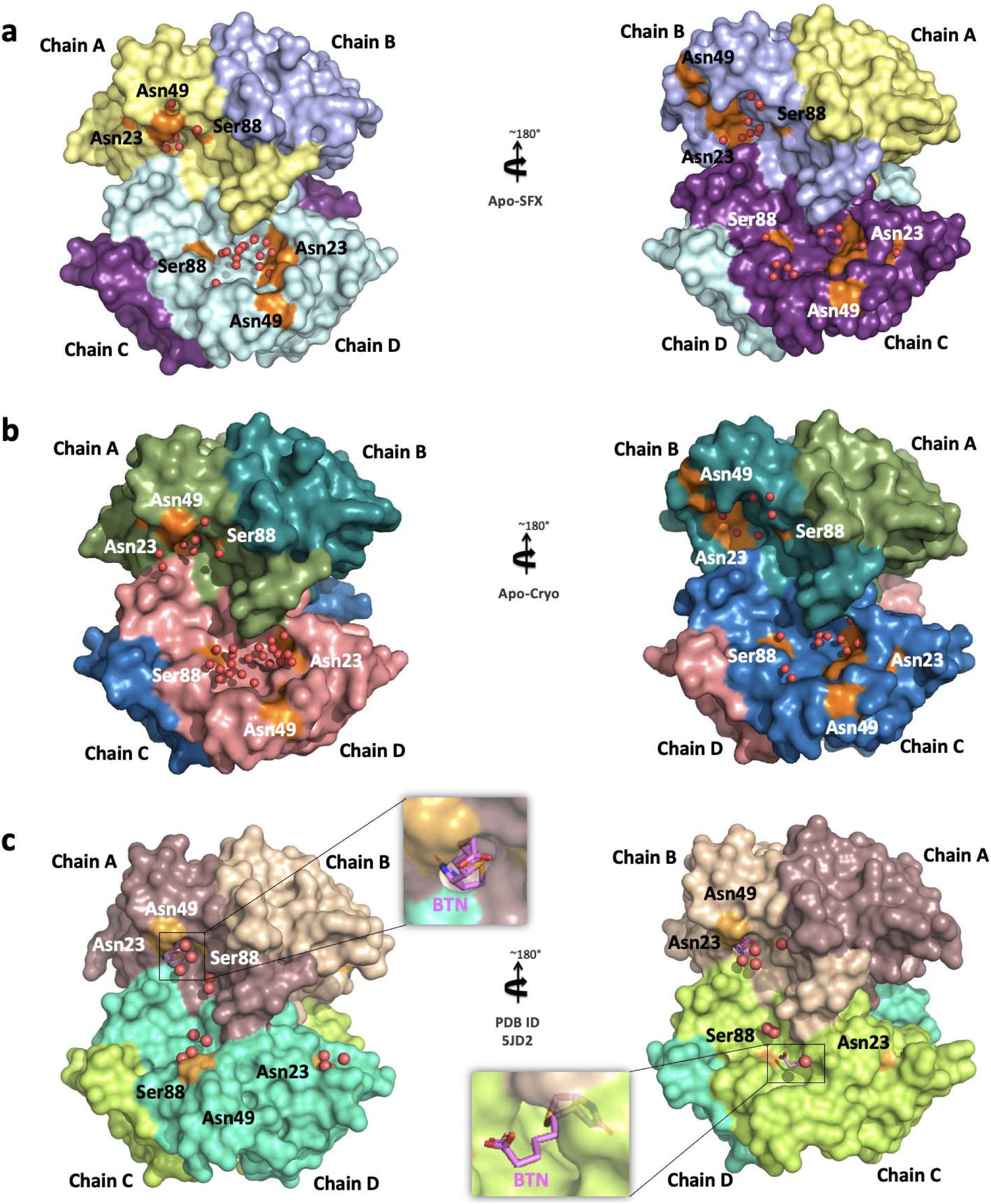
Representation of water molecules in the binding pocket of streptavidin structures. **(a)** Apo-SFX structure and **(b)** Apo-Cryo structure of apo-state streptavidin **(c)** Holo-SFX structure with selenobiotin (BTN) (PDB ID: 5JD2) are shown with their surface and water molecules were shown with red spheres. The indicated residues around the binding pocket are shown with orange color.

**Fig. 5:**
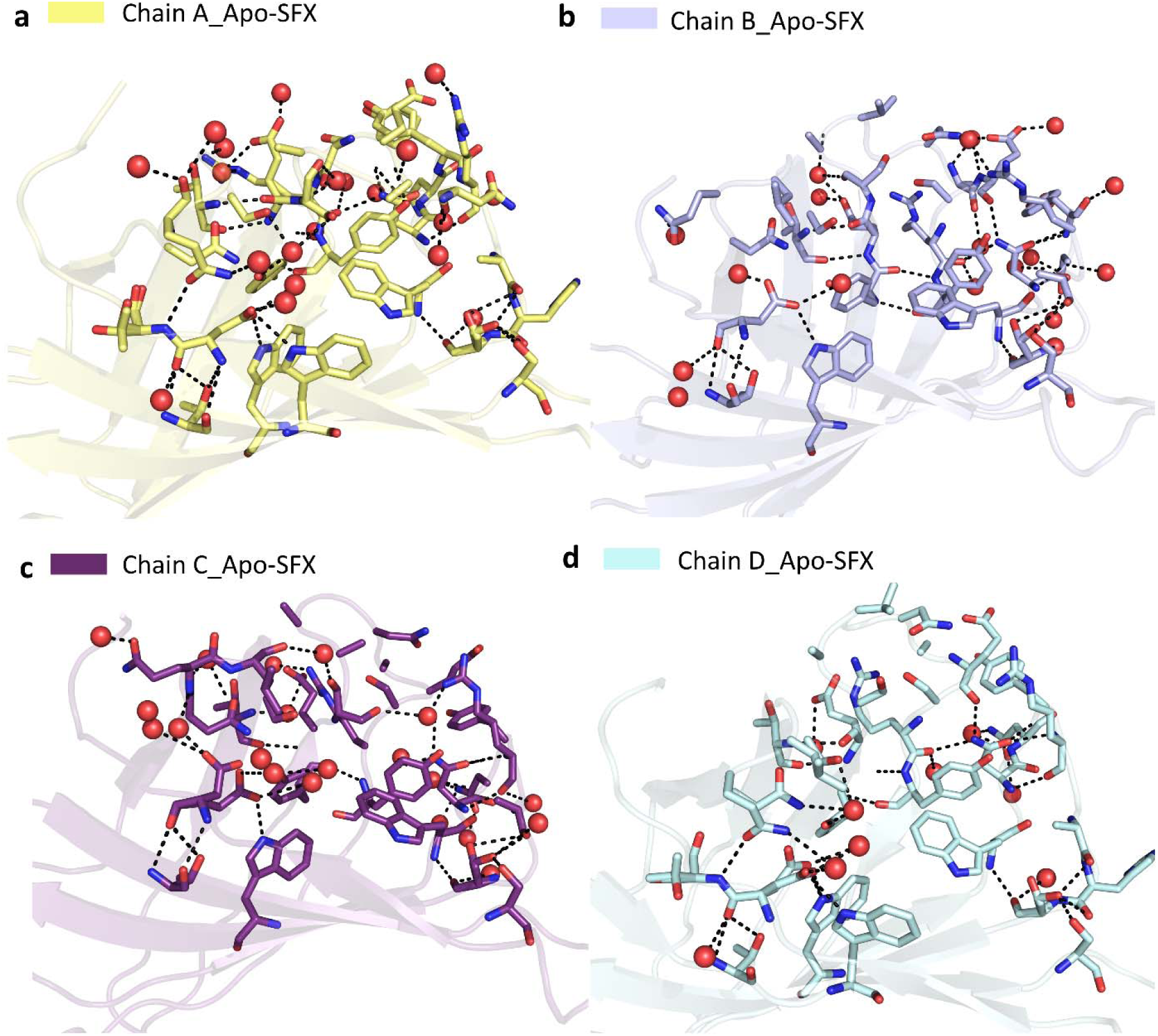
Representation of coordinated water molecules and polar interactions near the binding sites for each chain of Apo- SFX structure. Coordinated water molecules within the binding pocket were altered and polar interactions were reduced with loop opening in Apo-SFX structure. All polar interactions were observed within 3.6 Å**. (a)** Binding site residues of Chain A were observed with 19 water molecules and 51 polar interactions. Those interactions provided the stability of the chain which was similar to the selenobiotin bounded structure. **(b)** Chain B binding site residues were determined with 19 water molecules and 46 polar interactions that were involved. **(c)** Residues of the binding site of Chain C made 40 polar interactions between residues and 20 water molecules. **(d)** Chain D binding site residues involved 9 water molecules and 31 polar interactions. All interactions included H-bonds and electrostatic interactions and were presented with dashed lines. Water molecules were indicated with red-colored spheres.

To determine the structural dynamics of each subunit together, temperature factor analysis was performed and compared with the Holo-SFX structure **(Fig. 6a-c)**. While the two crystal structures of streptavidin reveal an inelastic binding pocket for chains A and B, we observed more intrinsic plasticity, especially in the loop region around the binding pocket of chains C and D. Electrostatic forces carry significant importance for protein-protein interactions and protein stability. The biotin-binding pocket for each chain was surrounded by a basic region concentrated with positively charged residues **(Fig. 7a-c)**. These surface charge redistributions can cause shape perturbations of a protein by altering hydrogen bonds and salt bridges.

**Fig. 6:**
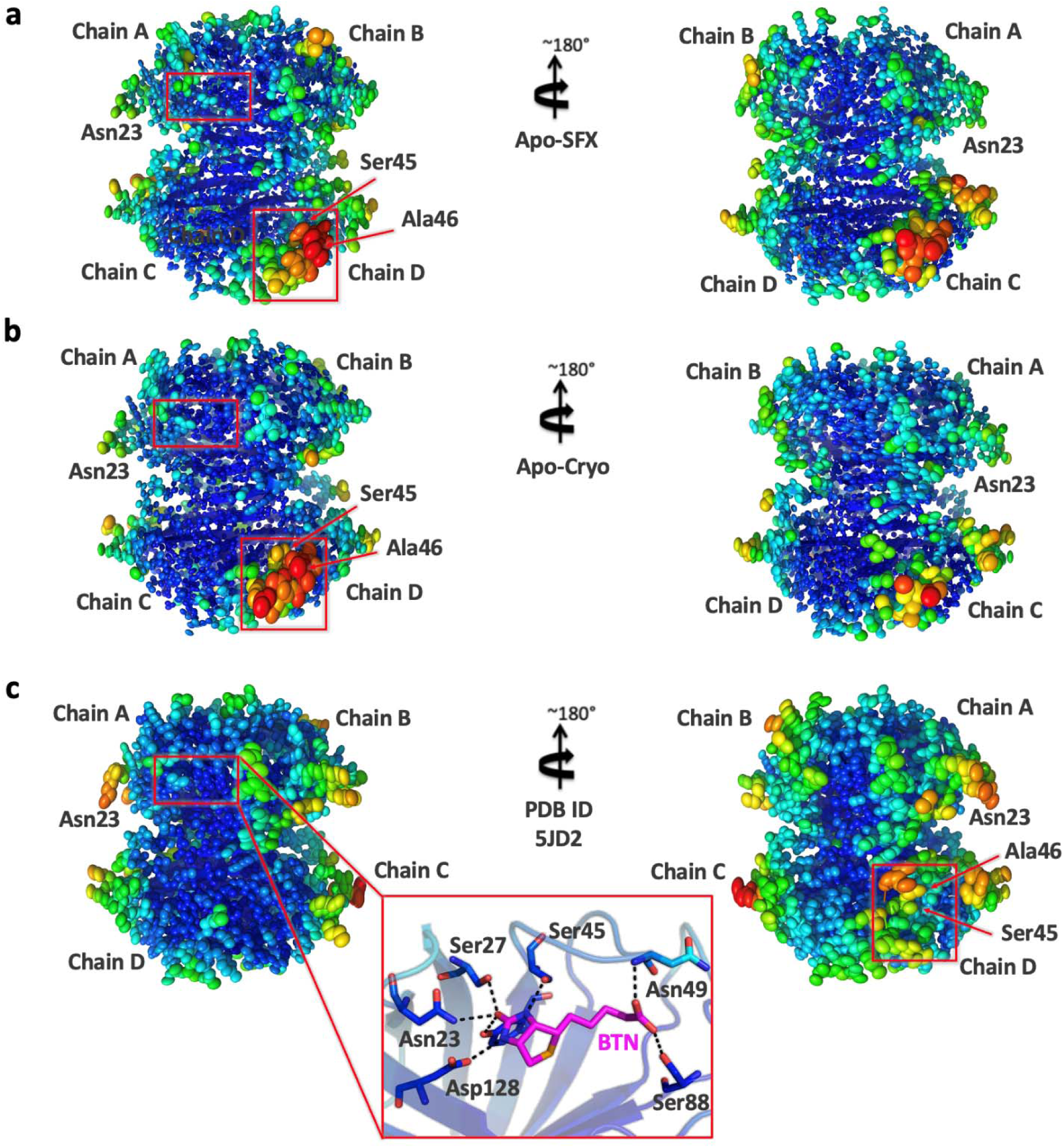
Representation of thermal ellipsoid structures of streptavidin structures. **(a)** Apo-SFX structure of streptavidin **(b)** Apo-Cryo structure of streptavidin **(c)** Holo-SFX streptavidin in complex with selenobiotin (PDB ID: 5JD2) are shown t determine the stability. Red boxes indicate flexible (red/orange) and stable (blue/green) regions on the streptavidin structures via b-factor presentation. The stable binding pocket of biotin is shown in panel C.

**Fig. 7:**
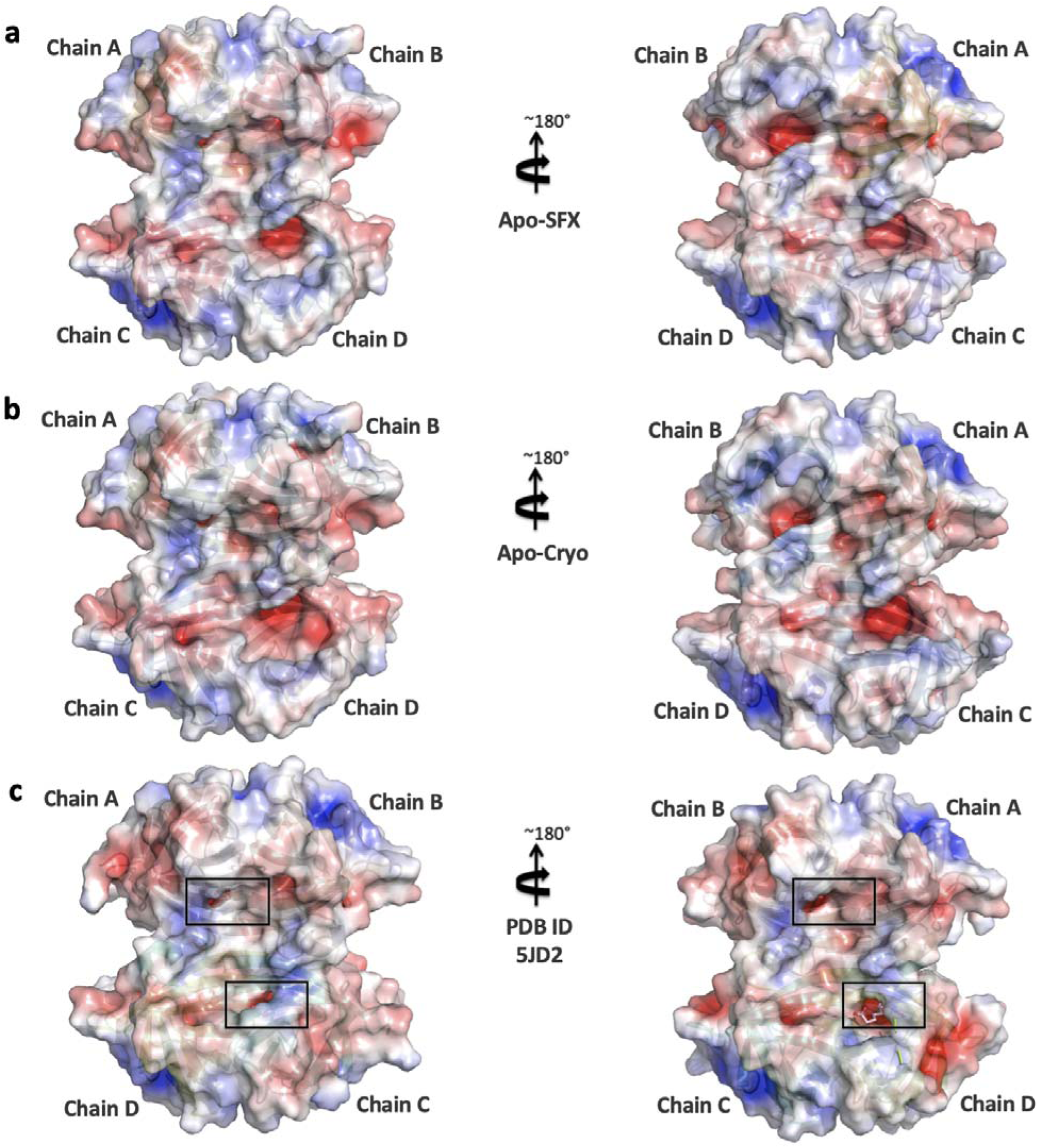
Representation of electrostatic surfaces. **(a)** Apo-SFX structure of streptavidin **(b)** Apo-Cryo structure of streptavidin **(c)** Holo-SFX streptavidin in complex with selenobiotin (PDB ID: 5JD2) are shown to detect the charge distribution. Holo-SFX structure is colored in light pink and shown with squares.

### Gaussian Network Model (GNM) Analysis

The GNM analysis is a coarse-grained Normal Mode Analysis method that is efficient in revealing a protein’s dynamics [36]. The normal modes obtained describe the microstates accessible to the protein’s native state. The theoretical fluctuations calculated with normal modes from GNM correlate with the thermal fluctuations found in X-ray experimentation as well. Slow modes have the highest mode weights and contain the most collective residue motions. These are the intrinsic fluctuations required for the protein’s global motion [37]. Therefore, Apo-SFX and Holo-SFX structures were analyzed with GNM to compare the apo- and holo-state streptavidin dynamics thoroughly. Cross-correlations between residue motions were investigated and mean squared fluctuations that describe the flexibility of the residues were analyzed. The theoretical fluctuations calculated from all modes were aligned to the B-factors in **Supplementary Fig. 14a,b**. They showed high correlation at the selected cut-off distance of 7.3 Å: overall correlation with B-factors was 0.785 in the GNM of Holo-SFX structure and 0.646 in the GNM of Apo-SFX structure. Cross-correlations between residue motions were investigated across all different modes in order to inspect the communication in the residue networks. Accordingly, chains A and B; C and D were observed as dimers with highly correlated motions in both Apo-SFX and Holo-SFX structures **(Fig. 8a,b)**. The active site residues of the Holo-SFX structure and selenobiotin ligand interactions of the related chains showed highly correlated motions as expected **(Fig. 8a)**. The intrachain residues’ cross-correlation section further showed that the first fifty residues had highly correlated motions. These residues belong to the same LJ-sheet with an extensive H-bond network. Moreover, residues after the 127th are highly correlated with those of the intrachain sections in both Apo-SFX and Holo-SFX structures **(Fig. 8a,b)**. Remarkably, correlated motion was especially high between 23-30th and 96-110th positions; 83-84th and 49-51st positions, respectively. Besides, residues in Trp120-Asp128 showed correlated motion with same numbered residues of the associated chain on the other dimer for both the Apo-SFX and Holo-SFX structures **(Fig. 8a, b)**. The result is expected due to Trp120 creating a water channel with Asp128 for the removal of waters from the streptavidin binding pocket, thus enhancing ligand binding [18]. The interchain cross-correlation sections within the dimers show highly correlated motions between residues Thr57 and Arg59; Thr71 and Lys80; His87 and Ser93; Leu110 and Gly113. Similarly, Asp61 has correlated motion between residues Thr76 and Thr90 of the neighboring chain on the same dimer **(Fig. 8a,b)**.

**Fig. 8:**
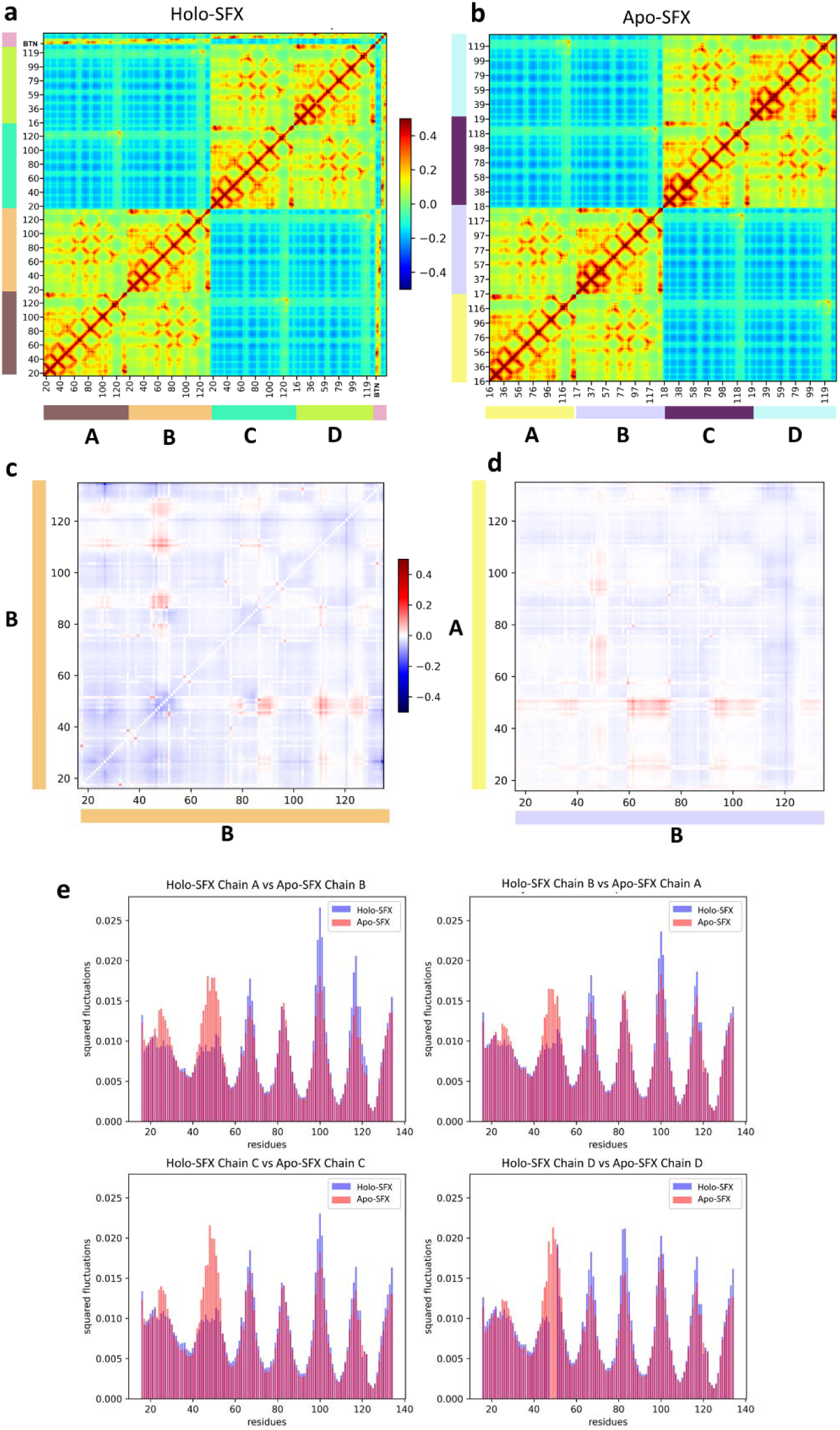
Gaussian Network Model (GNM) Analysis results for Apo-SFX and Holo-SFX (PDB ID: 5JD2) structures of streptavidin with selenobiotin. **a)** Cross-correlation heat-map from overall GNM modes for Holo-SFX structure. Selenobiotin was indicated with “BTN’’ in the figure **b)** Apo-SFX structure GNM analysis heat-map results from-overall modes. Highly correlated residue motion was represented with red and anti-correlated residue motions with blue color in A-B. **c)** The differences between intrachain cross-correlations of chain B for Holo-SFX structure over Apo-SFX structure results. **d)** Differences in the interchain cross-correlations of Holo-SFX structure over Apo-SFX structure at cross-sections of chains A and B. Differences in correlations were represented with blue color for decrease and red for increase **e)** Mean Squared Fluctuations of each chain of Holo-SFX structure and Apo-SFX structure was calculated from the 10 Slowest Mode from GNM analysis.

Cross-correlation differences between selected chains of Apo-SFX and Holo-SFX structures were also evaluated **(Fig. 8c,d)**. In these heat-maps, red indicates an increase in correlated motion between residues relative to the Apo-SFX structure, and blue means a decrease. Accordingly, when comparing the same chain of the two structures, residues between 45-51 have a higher correlation with residues 86-93, 110-114 and 121-129 within the Holo-SFX streptavidin than Apo-SFX streptavidin (**Fig. 8c**). Moreover, in the Holo-SFX structure, Ile17 and Val31, Ala35 and Ala38, Ser45 and Glu51, Thr57 and Arg59 displayed higher correlated motion when compared to Apo-SFX streptavidin. Furthermore, Apo-SFX streptavidin residues Gly26, Ala46 and Trp120 have higher correlation with all residues relative to the Holo-SFX structure (**Fig. 8c**). In particular, correlation of Gly26 with residues between 129-134 decreased for the Holo-SFX structure (**Fig. 8c**). On the other hand, correlation differences between the Apo-SFX and Holo-SFX structure of streptavidin are also different across chains in the same dimer (**Fig. 8d**). Interestingly, binding site residues have a higher correlation overall with residues of the opposite chain in Apo-SFX; correlation is highest with residues in position 16-40, 59-74 and 95-110 **(Fig. 8d**). In addition, Trp120 showed decreased correlation with residues of the opposite chain in the same dimer relative to Holo-SFX streptavidin (**Fig. 8d**).

Global protein motions were inspected closer to further determine the effects of selenobiotin binding on intrinsic residue fluctuations. Specifically, the mean squared fluctuations of the 10 slowest modes were obtained **(Fig. 8e).** These fluctuations reveal the protein’s global motion. Accordingly, fluctuation profiles of each chain demonstrate that regional flexibility contributes to the global motion of streptavidin. Thereafter, the Apo-SFX structure was observed with higher fluctuations around the selenobiotin binding site. Moreover, chains A to D showed increased flexibility, especially in the selenobiotin binding site, as evidenced by the high residue fluctuations. This result suggests that residues within L3/4, adjacent to the binding site, are intrinsically disordered and transitioning in between closed and open conformations in the absence of the ligand. Besides, presence of selenobiotin binding with streptavidin was observed with higher fluctuations around the 66, 110 and 116th residues. It must be noted that disorder was observed in chain D of the Holo-SFX structure at L3/4 **(Fig. 3d)** which also can be clearly seen from the GNM analysis graph **(Fig. 8e)**. Additionally, although the conformations should be similar in Apo-SFX chain A for both structures, fluctuation differences were observed between them at binding sites.

## DISCUSSION

The quaternary structures represent the highest complexity state in the structural hierarchy. The assembly of subunits interacting via noncovalent and covalent bonds determine their structure-function correlations [38]. Oligomeric proteins are presumed to be more robust and have evolutionary advantages over monomeric ones [39, 40]. The prevalence of proteins with a quaternary structure in biological systems is closely related to cooperativity [41]. There is no certain theoretical definition of this cooperativity in enzymology. The tetrameric structure of streptavidin is known as having the strongest binding affinity found in nature to its substrate biotin and it may thus further contribute to the understanding of cooperativity [18,34,42]. Simultaneous binding of four biotin ligands to streptavidin with full occupancy is unlikely [14]. Our previous structure of tetrameric Holo-SFX also displayed comparable properties with partial binding of selenobiotin to chain D [43]. In GNM analysis, the interchain residues displayed correlated motions with neighbour residues in the same dimers. Moreover, binding site residues generally have higher correlations with residues of the opposite chain (especially between 16-40th, 59-74th and 95-110th residues) in the Holo-SFX structure. In addition, Trp120 of the Apo-SFX structure had a higher correlation with overall residues than the Holo-SFX structure. This may confirm that the apo form of streptavidin displays tetrameric cooperativity via Trp120. Besides, biotin-binding stabilizes the L3/4 of streptavidin in tetrameric form which can be observed with lower fluctuations **(Fig. 8e)**. However, there are increased fluctuations such as Trp120 which are important for interchain connection within the tetramer **(Fig. 8e)**. Although a previous study indicated that the binding of biotin is not cooperative according to PCA analysis [44], these results confirm biotin binding may not have a stabilizing effect but can cause the allosteric response of streptavidin [23]. Also, biotin binding may affect other dimer chains of the tetrameric structure and provide cooperativity for additional ligand binding by perturbing active site residues.

In this study, we examined the polar interaction network of each subunit of the tetrameric Apo-SFX structure (**Fig. 5a-d**). We compared the binding sites of the Holo-SFX structure with the Apo-SFX structure (**Fig. 3a-d**). L3/4 helps ligand binding by removing excessive solvent which destabilizes H-bond interactions of the active site with a ligand. The free energy increase of 27.1 kcal/mol from the open to closed state transition of L3/4 was indicated [42]. It has been demonstrated that the open conformation of L3/4 is more favorable than the closed conformation [42]. This suggests that the “lid” closing is not spontaneous; however, the opening is spontaneous because of a lack of stabilization of an extensive polar interaction network with biotin. As a driving force, the closing of the lid may be caused by the allosteric and cooperative “pump-like” motions of tetrameric streptavidin which forcibly removes water molecules from the ligand binding site. This model is consistent with a previous study which indicated biotin dissociation is initiated by water entry and weakening H-bonds between binding site residues and biotin [18]. This may suggest cooperativity of a tetrameric form providing a higher affinity due to the removal of excess water molecules for establishing an extensive H-bond network in the binding pocket.

Open-loop conformation of the Apo-SFX streptavidin structure was supported with a B-factor analysis and represented as an ellipsoid model [45] (**Fig. 6a-c**). Based on GNM results, the L3/4 was observed with higher fluctuations in Apo-SFX streptavidin, representing higher mobility of the residues **(Fig. 8e)**. One reason for the striking difference in fluctuations is that selenobiotin was included in the GNM of Holo-SFX (5JD2) and, therefore, the ligand’s effect on decreasing the flexibility of the region was more clear in the protein’s global motions. The tetrameric form of the enzyme is the most efficient form of the streptavidin-biotin complex when compared to dimeric and monomeric forms [14]. Interestingly, we observed higher correlated motions of residues in dimers which may indicate dimeric cooperativity of the streptavidin **(Fig. 8a,b)**.

Chain A of our Apo-SFX structure was captured in a closed-loop conformation with a continuous electron density of water molecules suggesting that those water molecules may be mimicking partially occupied biotin **(Supplementary Fig. 15a)**. This is consistent with a previous study that had comparable results at chain A [46]. On the other hand, streptavidin-avidin affinity comes from a hydrophobic cage caused by a Van der Waals and hydrogen bond network associated with water molecules [15] (**Fig. 5a-d and Fig. 7a-c**). The intrinsic plasticity of L3/4 via cooperative allosteric effects of these residues may be the cause of an asymmetrical binding event of biotin. The polar interaction networks and number of water molecules in contact with streptavidin are directly related to the L3/4 conformation. It has a closed conformation through increased polar interactions at the binding site **(Fig. 5a-d & Supplementary Fig. 15a)**. However, decreases in coordinated water molecules are associated with an open conformation of the loop, acting as a “lid” **(Fig. 4a-c & Supplementary Fig. 15b-d)**. H-bond formation is a dynamic process [47] and these alternative extensive polar interaction networks of tetrameric streptavidin may contribute to the observed asymmetry. Similar to the Holo-SFX structure **(Fig. 3a-d)**, Holo-CryoEM structure of streptavidin in complex with biotin (PDB ID: 6J6J) was superposed with our Apo-SFX structure **(Supplementary Fig. 9a-d).** The comparison of these structures suggests that the two techniques can capture alternative binding conformations and expand the conformational space sampling of the active site loop.

Previous studies described that L3/4 may or may not be randomly open in three monomers and closed in the last one. This open/closed loop conformation cannot be caused by crystal-packing interactions [46, 48]. Our data suggest that this structural feature is intermolecular allostery between monomers of streptavidin. Recently, a new model was proposed to describe structural cooperativity of streptavidin and a cooperative allosterism upon binding biotin [23]. Ligand binding is characterized by L3/4. It functions as a “lid” that closes the binding pocket when it binds to biotin. In the Apo-SFX structure, L3/4 appears very flexible and unmodeled. However, our atomic-resolution structural data and GNM analysis suggest that there is a predisposed streptavidin cooperative allosterism not mediated by the first biotin molecule. Moreover, the open conformation of L3/4 is observed in three subunits while the fourth L3/4 region is observed in closed-state, which is a harmonic motion that triggers the first biotin binding **(Supplementary Fig. 16a-d & Supplementary Fig. 17)**. The tetrameric structure of streptavidin provides higher affinity by water channels and “lid” like L3/4 movement caused by those cooperative and allosteric binding events. To determine and validate the accuracy of previous structures of streptavidin, the SFX approach offers structural data without temperature or radiation damage, providing a solid template for future studies. The next step to better understand the details of this binding and cooperative allosterism is performing time-resolved structural analysis by using ultrabright and ultrafast XFELs [49].

## MATERIALS AND METHODS

### Sample preparation for SFX crystallography and cryo-synchrotron crystallography

Purchased core-streptavidin protein (Cat# Streptavidin-501; Creative Biomart, USA) was added into DTT solution with a 1 mM final concentration. Then, streptavidin proteins were crystallized by sitting-drop microbatch screening under oil and using 72 well Terasaki crystallization plates. For crystallization, the protein was mixed with ∼3500 commercially available sparse matrix crystallization screening conditions (1:1 ratio, v/v) at ambient temperature. The best crystals were grown in Pact PremierTM 100 mM MMT buffer pH 6.0 and 25 % w/v PEG 1500 and were covered with 100% paraffin oil. Microcrystals 1-5 × 5-10 × 10-20 μm^3^ in size were passed through 100 micron plastic mesh filters (Millipore, USA) in the same mother liquor composition to eliminate the large single crystals and other impurities before the data collection. Crystal concentration was detected as 10^10^-10^11^ particles per ml for SFX based on light microscopy. Initial crystals of the batched crystalline slurry were not pretested for diffraction quality before XFEL and synchrotron beamtime. For a better comparison with GNM analysis, Holo-SFX crystals and Apo-SFX crystals were obtained from the same batch with minimized variables by using the same crystallization conditions, mother liquor and protein sample.

### Transport of microcrystals for SFX studies at MFX instrument at the LCLS

Crystal slurry at the total volume of 1.9 ml was transferred to a 2 ml screw-top cryovial (Wuxi NEST biotechnology, China cat#607001). These vials were wrapped loosely by Kimwipes (Kimberly-Clark, USA) and tightly placed in 20 ml screw top glass vials to provide insulation and prevent mechanical shocks during air transportation. The vials were wrapped with excess amounts of cotton (Ipek, Turkey) and placed in a ZiplocTM bag (SC Johnson, USA) to provide both added layer of insulation and mechanical shock absorption. Furthermore, ZiplocTM bags are placed into a styrofoam box which was padded with ∼1 kg of cotton and covered with an additional layer of 1 cm thick loose cotton. Packing the samples with cotton, which prevented physical damage of the crystals, and the successful transportation to XFEL, was followed by diffraction to 1.7 Å resolution.

### MESH sample injection of streptavidin crystals for SFX studies at MFX instrument at the LCLS

The 1.6 ml sample reservoir was loaded with streptavidin crystal slurry in their unaltered mother liquor as described above. The Microfluidic Electrokinetic Sample Holder (MESH) injector was used for the injection of streptavidin crystals [50]. The sample capillary was a 200 μm ID × 360 μ OD × 1.0 m long fused silica capillary. The applied voltage on the sample liquid was typically 2500-3000 V, and the counter electrode was grounded. The sample flow rate was typically between 2.5 and 8 μl/min.

### Data collection and analysis for SFX studies at LCLS

SFX data was collected during the LCLS beamtime (ID: mfxp17318) at the MFX instrument of LCLS at SLAC National Accelerator Laboratory (Menlo Park, CA). The radiation damaged-free diffraction data collected from streptavidin microcrystals on a ePix10k2M pixel array detector installed at the MFX instrument. X-ray beam with a vertically polarized pulse with duration of 30 fs was focused using compound refractive beryllium lenses to a beam size of ∼6 × 6 μm full width at half maximum (FWHM) at a pulse energy of 0.8 mJ, a photon energy of 9.8 keV (1.25 Å) and a repetition rate of 120 Hz. OM monitor [51] and *PSOCAKE* [52, 53] were used to monitor crystal hit rates, analyze the gain switching modes and determine the initial diffraction geometry of the detector [54]. The detector distance was arranged as 18 mm, with an achievable resolution of 2.1 Å at the edge of the detector and 1.64 Å at the corner of the detector. A total of 691,200 detector frames were collected continuously without any clogging issues from streptavidin microcrystals.

### Data collection and analysis for cryo-synchrotron studies at SSRL

Synchrotron X-ray diffraction data were collected from a single crystal for apo-streptavidin with a Pilatus 6M detector at microfocus beamline BL12-2 at the Stanford Synchrotron Radiation Lightsource (SSRL) in Menlo Park, CA. The diffraction data in space group P21 were collected to 1.1 Å resolution with unit cell dimensions a=46.36 Å b= 85.76 Å c=58.13 Å α 0.00 β 8.71 γ=90.00 at a wavelength of 0.979 Å and -180 °C.

### Data processing for SFX and cryo-synchrotron structures; hit finding, indexing and scaling

The diffraction data for the SFX structure were collected through the MFX instrument using ePix10k2M detector. Total diffraction patterns were selected as potential crystal hits using *CHEETAH* software [55]. The hit finding, which is based on Bragg reflections, was performed by using peakfinder8 and the images containing more than 20 peaks were classified as crystal hits that were indexed by using the *CrystFEL* software package [56, 57] version 9.0 [58]. While *XGANDALF* [59], *DIRAX* [60], *MOSFLM* [61] and *XDS* [62] were used as indexing algorithms, the indexed reflections were subsequently integrated and merged using *PARTIALATOR* [63] applying the unity model over 3 iterations and the max-ADU set to 7500. The complete reflection intensity list from *CrystFEL* was then scaled and cut using the *TRUNCATE* program from the *CCP4* suite [64] prior to further processing. For streptavidin crystals, the final dataset included 384,250 hits with a total of 106,021 indexed patterns ( 28%) was merged into a final dataset (P12_1_1, unit cell: a = 47.40 Å, b = 87.70 Å, c = 58.90 Å; α = 90.00, β = 98.90, γ = 90.00) **(Table 1)**. Additionally, using a resolution cutoff at 1.7 Å, an R_split_ of 11.0% was obtained along with a CC* of 0.99 over the entire resolution range. The final dataset had an R_split_ of 77.8%, and CC* of 0.87 in the highest resolution shell. For cryo-synchrotron structure, X-ray diffraction data (P12_1_1, unit cell: a = 46.36 Å, b = 85.76 Å, c = 58.13 Å; α = 90.00, β = 98.71, γ = 90.00),were collected by using a Dectris Pilatus 6M detector installed at BL-12-2 instrument at SSRL, which was processed with *XDS* [62] package for indexing and scaled by using *XSCALE* [62]. The resolution cutoff set to 1.1 Å without negatively impacting R_free_ and R_work_ **(Table 1)**.

### Structure determination and refinement of crystal structures

Apo-streptavidin structures, obtained at LCLS (MFX) and cryo-synchrotron at SSRL (BL12-2), were determined by using the automated molecular replacement program *PHASER* [65] implemented in *PHENIX* software [66] with the previously published Holo-SFX structure as a search model and initial rigid-body refinement [43]. After a simulated-annealing refinement, individual coordinates and TLS parameters were refined. Potential positions of altered side chains and water molecules were checked by using the program *COOT* [67]. Then, positions with strong difference density were retained. The Ramachandran statistics for Apo-SFX structure (most favored/additionally allowed/disallowed) were 98.31/1.48/0.21% respectively. The Ramachandran statistics for cryo-synchrotron structure (most favored / additionally allowed / disallowed) were 97.47/2.11/0.42 % respectively. For an understanding ligand binding dynamics, Apo-SFX structure is aligned with the ambient temperature Holo-SFX structure [43] and figures were generated by *PyMOL* software [68].

### Temperature factor analysis and generation of ellipsoids

Both SFX and cryo-synchrotron structures were examined to generate ellipsoid structures based on B-factor. The generation of ellipsoid models via *PyMOL*[68] was enabled based on structure refinement with TLS parameters through *PHENIX* [69]. Later on, these two structures were compared with the Holo-SFX structure to provide better understanding of the flexibility of atoms, side chains and domains. The all ellipsoid structures were colored with rainbow selection on *PyMOL* [68].

### Gaussian Network Model (GNM) Analysis

Apo-SFX streptavidin and Holo-SFX streptavidin were analysed by Normal Mode Analysis with Gaussian Network Model (GNM) using *ProDy* [70]. Contact maps were defined with all Cα atoms of the proteins (residue numbers between 15-135). The selenobiotin atoms N1, C2, C9 and O12 were included in the holo structure’s GNM. Thus, the Apo-SFX streptavidin had 476 atoms selected to build a Kirchhoff matrix and the selenobiotin bound structure had 486. Same cutoff distance of 7.3 Å was selected in both models to assume pairwise interactions. Default spring constant of 1.0 was used for both structures. All normal modes were calculated with GNM: 475 non-zero modes were obtained for the apo structure and 488 non-zero modes for the holo structure. The theoretical fluctuations calculated with all GNM modes were compared with the experimental B-factors. Cross-correlations between residue fluctuations were determined over all GNM modes as well. The differences in cross-correlations between Holo-SFX and Apo-SFX structures were calculated at selected sections: intrachain cross-correlations of chain A in Apo-SFX were subtracted from the intrachain cross-correlations of chain B in selenobiotin bound structure; and the interchain cross-correlations between chains A and B were subtracted similarly. Their results were presented as heat-maps. The selections were decided on after superposing the Apo-SFX and Holo-SFX structures: chain B of the apo-structure aligned with the chain A of the holo-structure. The 10 slowest modes were chosen for analysis of the global motions based on their high variance calculated with *ProDy*. The weighted squared fluctuations of the two structures at these modes were aligned to compare the selenobiotin bound and unbound state dynamics of streptavidin.

### Data Availability

Presented two apo-streptavidin in this article are available as 7EK8 and 7EK9 on the Protein Data Bank. Any remaining information can be obtained from the corresponding author upon reasonable request.

## Acknowledgments

The authors gratefully acknowledge use of the services and facilities of the Koç University IsBank Infectious Disease Center (KUIS-CID). HD acknowledges support from National Science Foundation (NSF) Science and Technology Centers grant NSF-1231306 (Biology with X-ray Lasers, BioXFEL). HD would like to thank Michelle Young, Ritu Khurana, Lori Anne Love and Tracy Chou for their invaluable support and discussions. This publication has been produced benefiting from the 2232 International Fellowship for Outstanding Researchers Program of TÜBİTAK (Project No: 118C270). However, the entire responsibility of the publication belongs to the owner of the publication. The financial support received from TÜBİTAK does not mean that the content of the publication is approved in a scientific sense by TÜBİTAK. Use of the Linac Coherent Light Source (LCLS), SLAC National Accelerator Laboratory, is supported by the U.S. Department of Energy, Office of Science, Office of Basic Energy Sciences under Contract No. DE-AC02-76SF00515. CD acknowledges support from TÜBİTAK (Project No: 120Z594). The authors gratefully acknowledge use of the services and facilities of the Koç University Research Center for Translational Medicine (KUTTAM), funded by the Presidency of Turkey, Presidency of Strategy and Budget. The content is solely the responsibility of the authors and does not necessarily represent the official views of the Presidency of Strategy and Budget. Portions of this research were carried out at Stanford Synchrotron Light Source (SSRL) at the SLAC National Accelerator Laboratory. SSRL is supported by the U.S. Department of Energy (DOE), Office of Science, Office of Basic Energy Sciences (OBES) under Contract No. DE-AC02-76SF00515. The SSRL Structural Molecular Biology Program is supported by the DOE Office of Biological and Environmental Research and by the National Institutes of Health, National Institute of General Medical Sciences (NIGMS) (including P41GM103393).

## Author contributions

HD, CD, FBE, GY prepared the samples. AB, ZS, FP, CHY, CK, RGS performed the sample delivery. On-site data collection. HD, CD, EA, BY, ED, FBE, GY, SB, EHD, BH, ML, MHS, MSH, AB, VM, ZS, FP, CK, AC, TD, RGS performed the remote data collection. SB, OMY, AB, AT, GKK, ZS, FP and CHY executed the data processing. Structures were refined by HD. Data analyzed by HD, EA, BY, ED. The manuscript was prepared by HD with input from all the coauthors.

---

**Supplementary Fig. 1:**
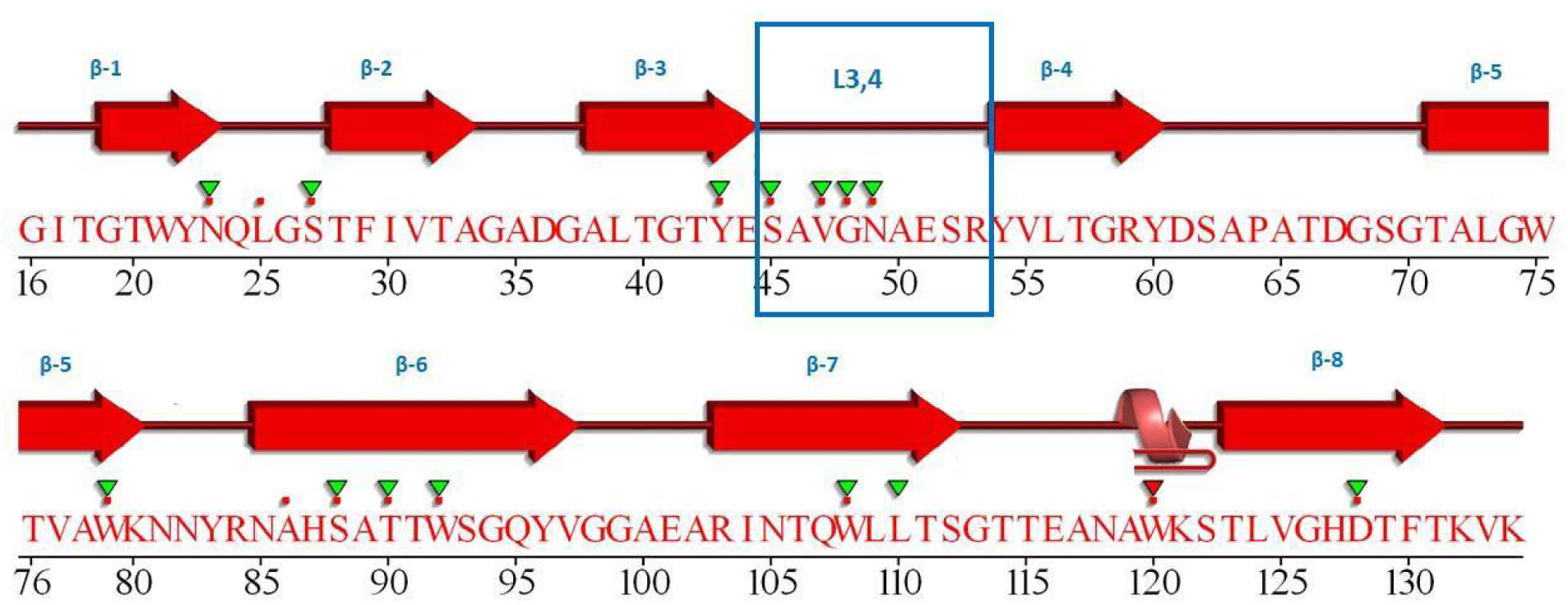
Secondary structure and sequence of streptavidin. The arrows indicate ß-strands, and the helix shape represents α-helice. Flat U turns shapes represent flexible loop. Loop 3/4 marked with a square and labeled as L3/4. Red dots represent residues in contact with ligand (selenobiotin), while inverted triangles that are colored in green and red represent functional residues of repeats. This figure was created using the PDBsum server and modified [71].

**Supplementary Fig. 2:**
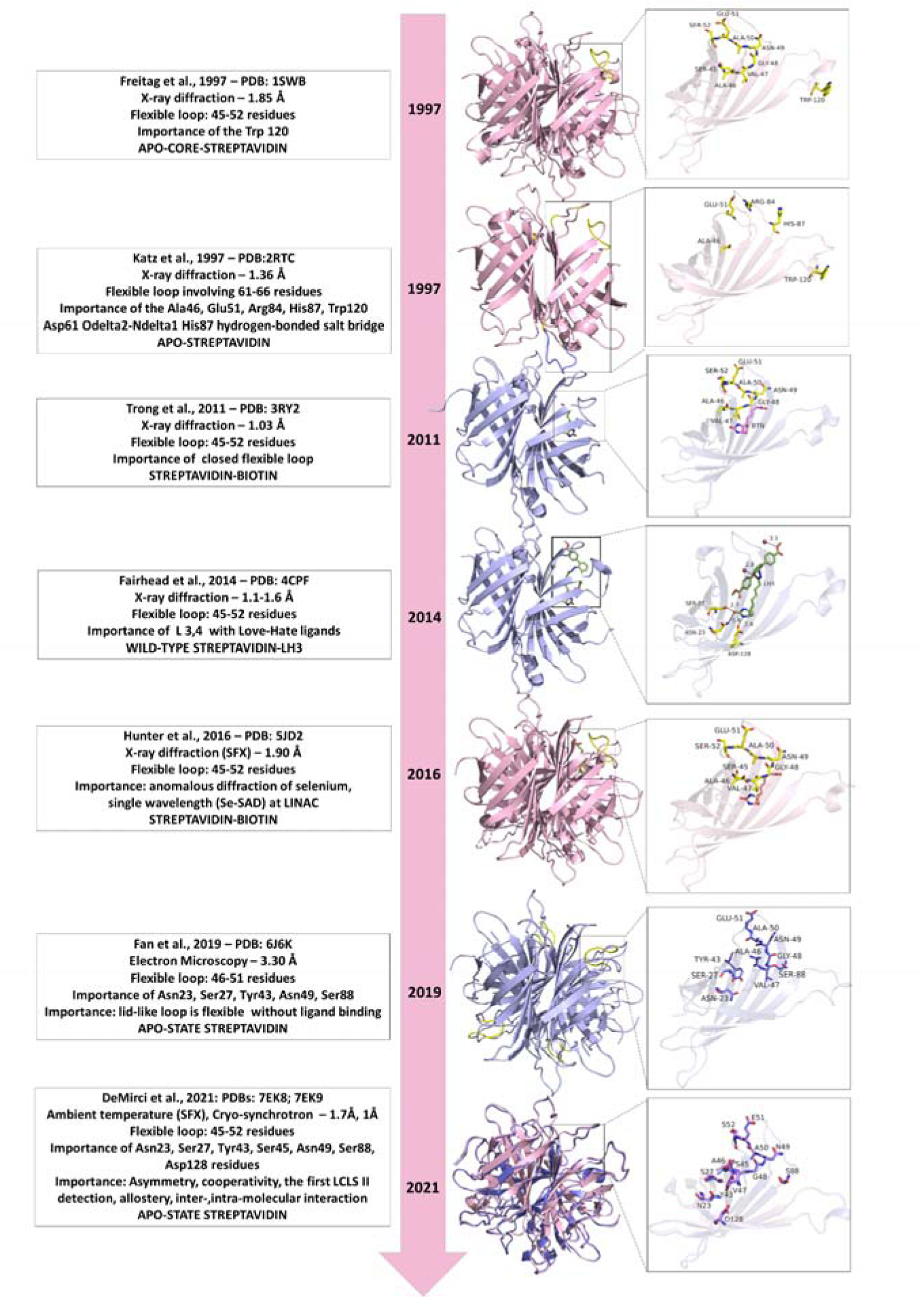
Chronological partial-bibliography of the apo- and holo-streptavidin structures. **1997:** It was emphasize on the 45-52 inter-residual loop (L3/4) concept and the importance of the Trp120 residue in apo-streptavidin (PDB_ID: 1SWB). **1997**: Simultaneous X-ray diffraction model. Its resolution is higher than the previous model. It was emphasized the importance of the loop 61-66 rather than L3/4 as well as of His87, Ala46, Glu51, Trp120and Arg-84. Additionally, it was indicated the importance of the salt bridge between Asp61 and His87 (PDB_ID: 2RTC). **2011:** Relatively high-resolution streptavidin-biotin model compared to the previous structures. Attention had been drawn to the concept of a “closed-flexible” L3/4 residues (PDB_ID: 3RY2). **2014:** In order to understand the protein plasticity, conflicted ligands were used instead of biotin. The importance of Ser-45 in L3/4 cycle is emphasized with the LH3-bound wild-type streptavidin structure (PDB_ID: 4CPF) **2016**: It was indicated the first SFX streptavidin-biotin model that has been studied with ambient-temperature X-FEL. The L3/4 conformation is pointed out, and for the first time, selenobiotinyl-streptavidin structure demonstrated by using phases obtained by the anomalous diffraction of selenium measured at a single wavelength (Se-SAD) at the Linac Coherent Light Source (PDB_ID: 5JD2). **2019**: It was demonstrated the first apo-state streptavidin model powered by Cryo-EM. Current work refers to the concept of a “lid” like loop without ligand-binding, paying attention to loop conformation between 51-56 residues and its interaction with active residues (PDB_ID: 6J6K). **2021:** This work emphasizes the first models to be extensively investigated apo-core streptavidin powered by radiation-damage free SFX (Apo-SFX, PDB: 7EK8) as well as high-resolution cryo-synchrotron (Apo-Cryo, PDB: 7EK9). The L3/4 conformation has been given importance. First LCLS II study, involving inter- and intra-monomeric examination, compares the interaction of its active residues with the previous study (PDB_ID: 5JD2).

**Supplementary Fig. 3:**
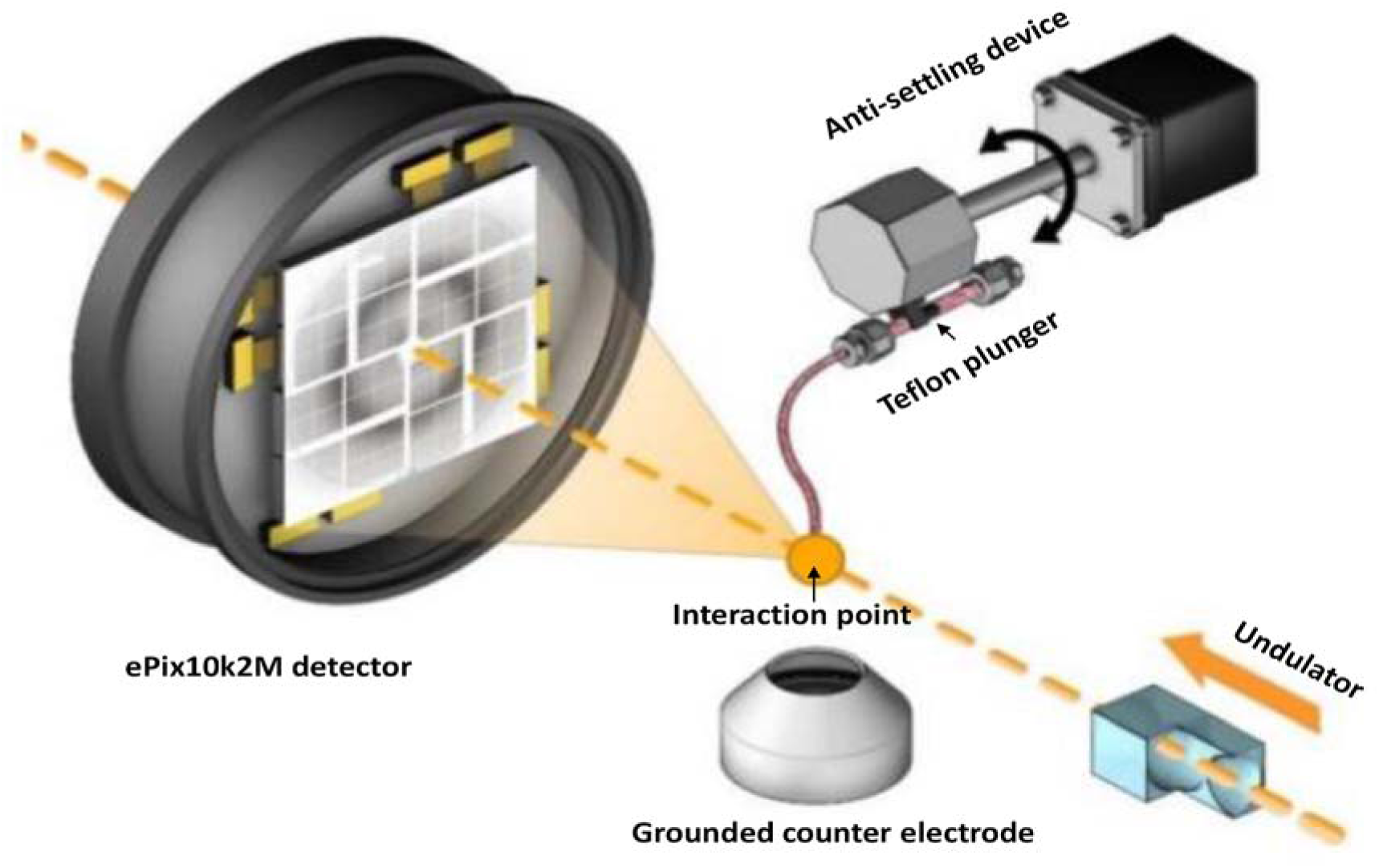
Diagram of the MESH injector setup at the XFEL. Protein microcrystals are injected by this injector. The sample reservoir has a Teflon plunger (indicated by arrow). This reservoir is attached to an anti-settling device rotating at an angle to prevent crystal settling of proteins and keep them homogenized. The protein crystals and the LCLS pulses interacted at the point indicated by the arrow. After the X-ray beam hits these crystals at the interaction point (indicated by an arrow), diffraction data is collected by the ePix10k2M camera.

**Supplementary Fig. 4:**
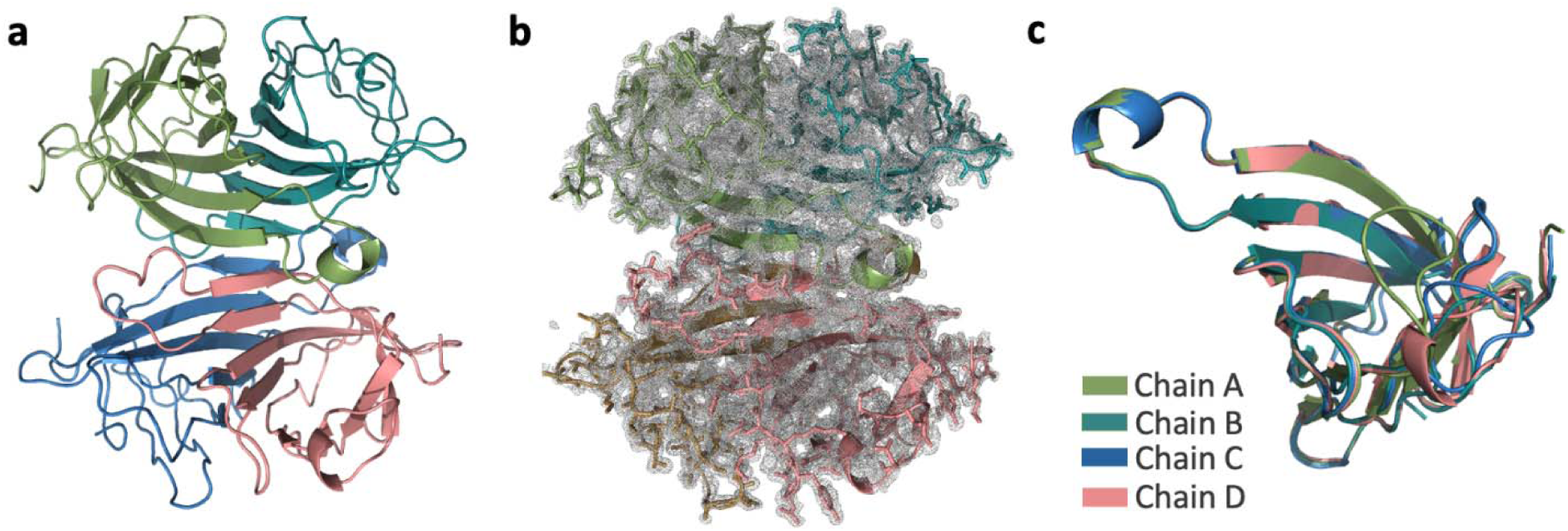
Synchrotron structure of streptavidin (Apo-Cryo). **(a)** The apo structure of streptavidin is colored based on each chain. **(b)** 2*F*o-*F*c simulated annealing-omit map at 1 sigma level is colored in gray. **(c)** Each chain of streptavidin is superposed with an overall RMSD of 0.136 Å.

**Supplementary Fig. 5:**
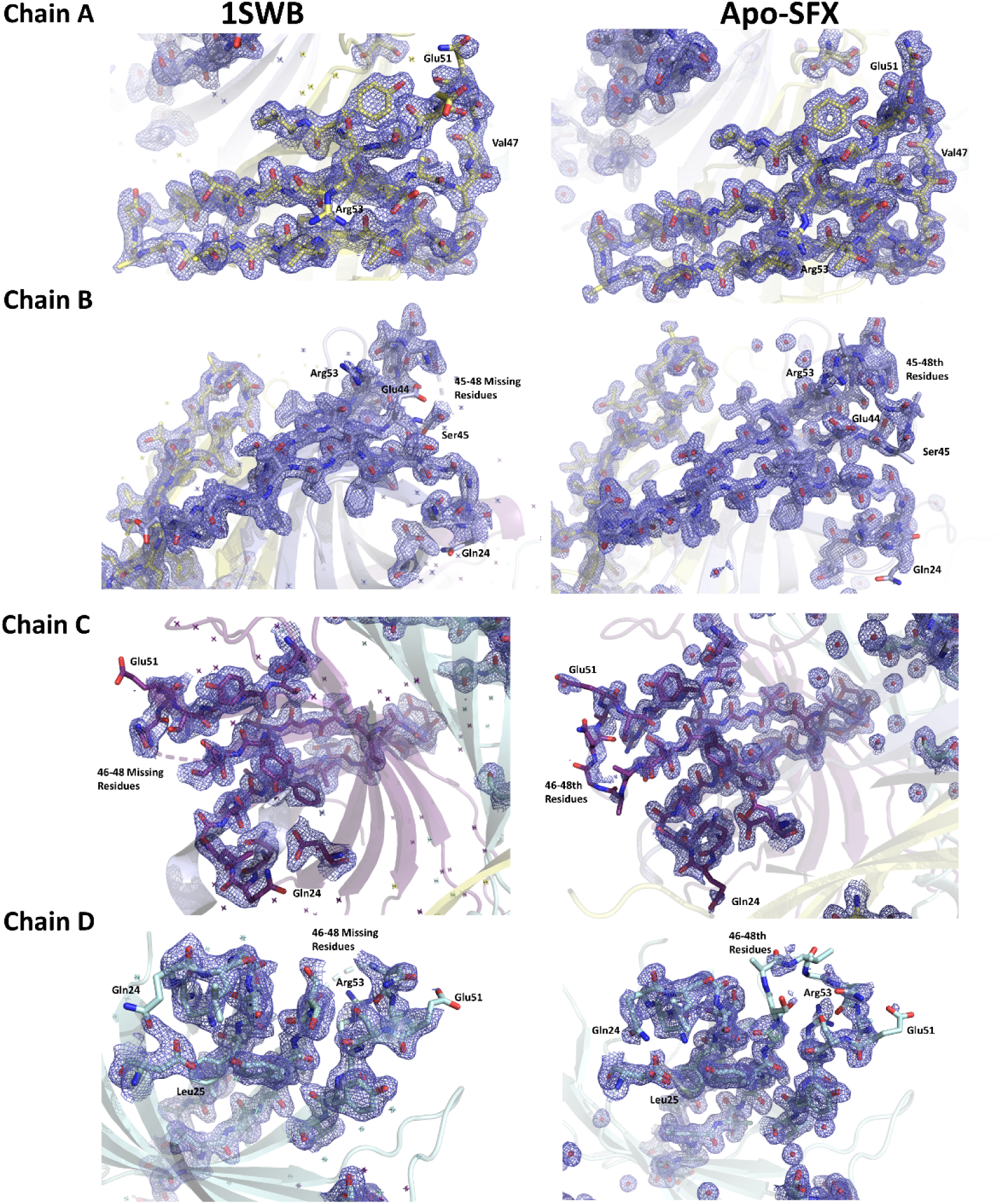
Electron density map comparison of binding site residues of the 1SWB and Apo-SFX structure of streptavidin. 2*F*o-*F*c simulated electron density map is colored in slate. Chain colors are presented as described before. 1SWB structure has missing residues between 46-48th position in chain B, C and D. Moreover, Gln24, Lue25, Val47, Glu51, Arg53 were observed with better electron density for Apo-SFX structure.

**Supplementary Fig. 6:**
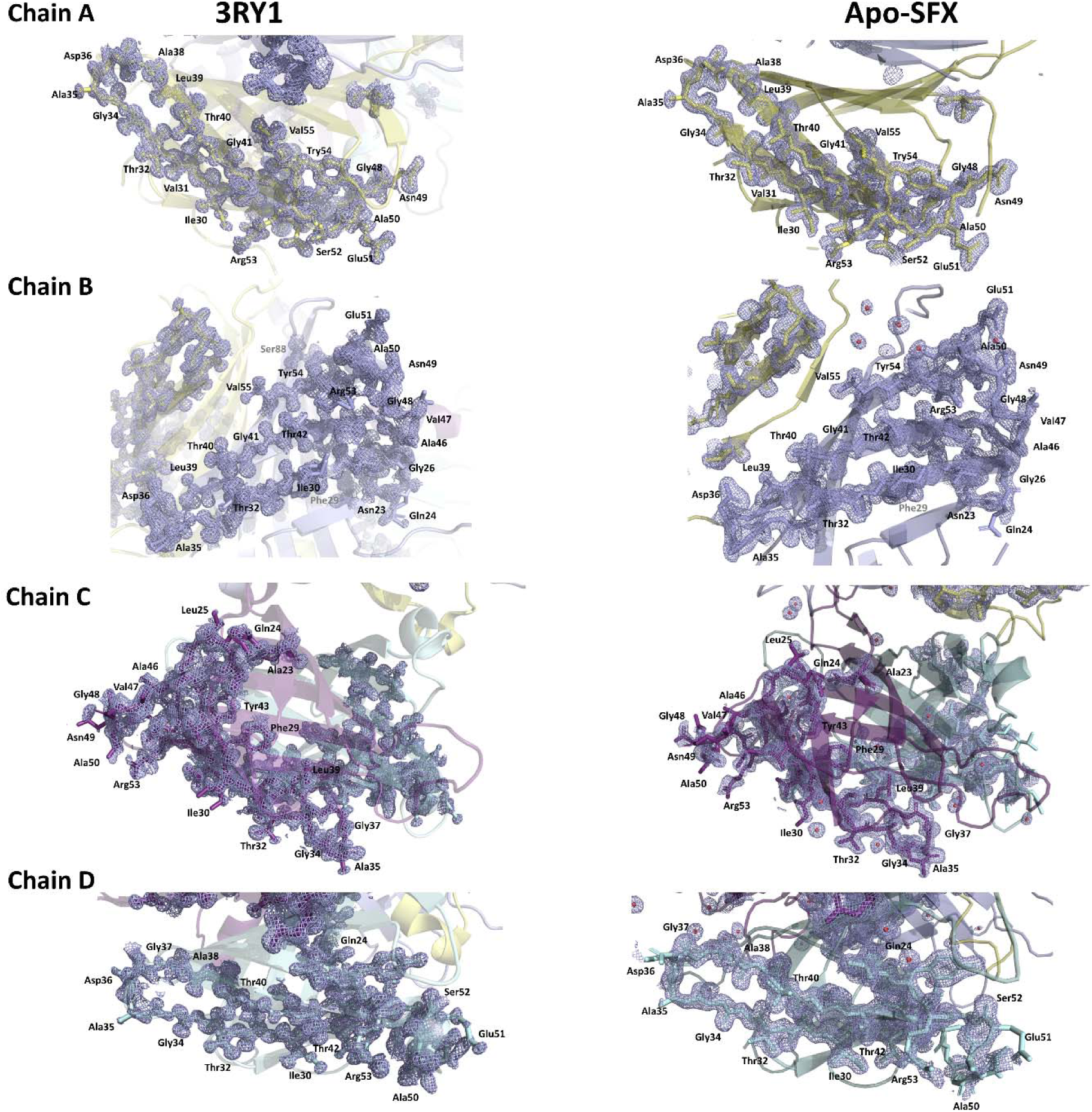
Electron density map comparison of binding site residues of the 3RY1 and Apo-SFX structure of streptavidin. 2*F*o-*F*c simulated electron density map is colored in slate. Chain colors are presented as described before. For Apo-SFX structure, chains A and B were observed with enhanced and continuous electron density for those binding residues compared to 3RY1 structure. However, in chain C and D, Apo-SFX structure was observed with better electron density at the beginning and end of the binding site residues such as Asn23 or Glu51, while L3/4 was identified with better electron density at 3RY1 structure.

**Supplementary Fig. 7:**
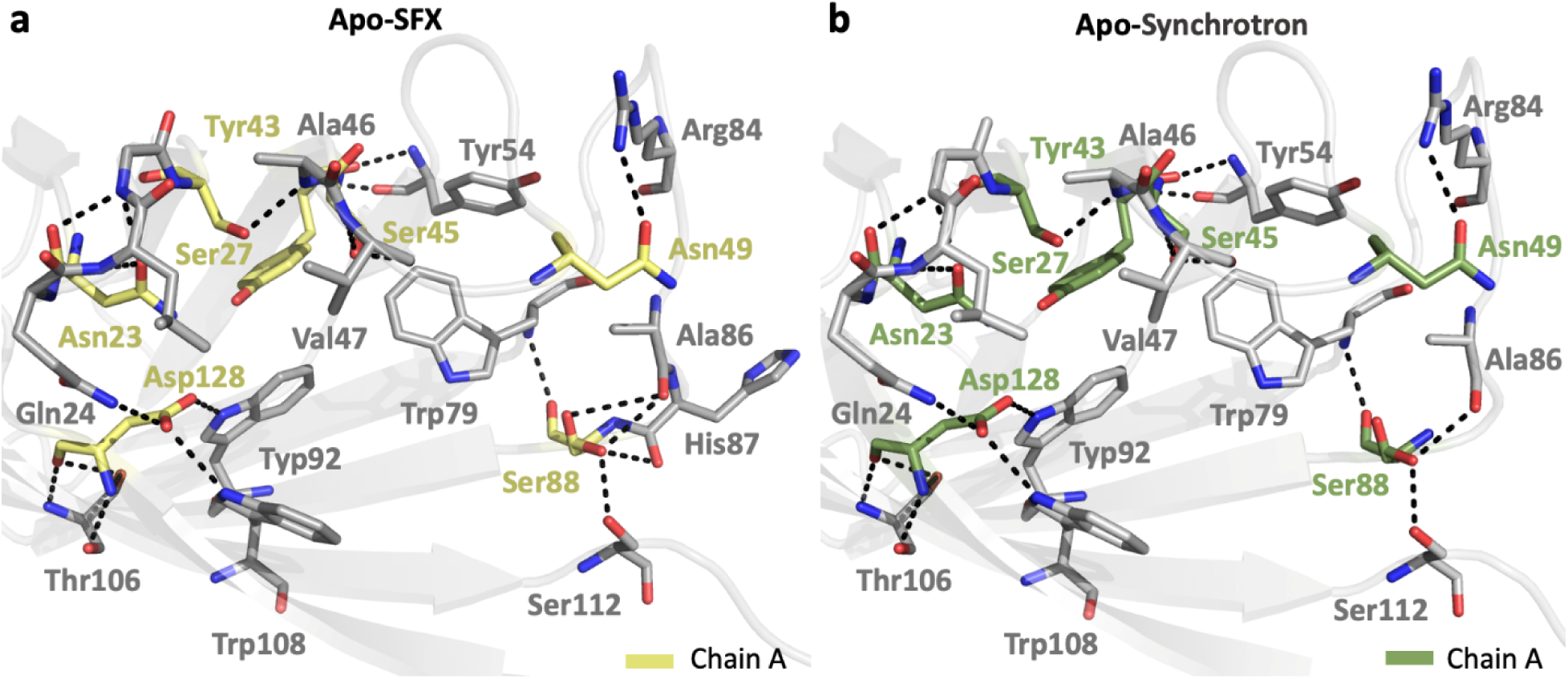
Hydrogen bond interactions in the binding pocket. **(a)** The interactions within 1-3.5 Å are represented by dashed lines and active residues in chain A of SFX structure are colored in pale yellow while flank cavity residues are colored in gray. **(b)** The interactions within 1-3.5 Å are represented by dashed lines and active residues in chain A of synchrotron structure are colored in green while flank cavity residues are colored in gray.

**Supplementary Fig. 8:**
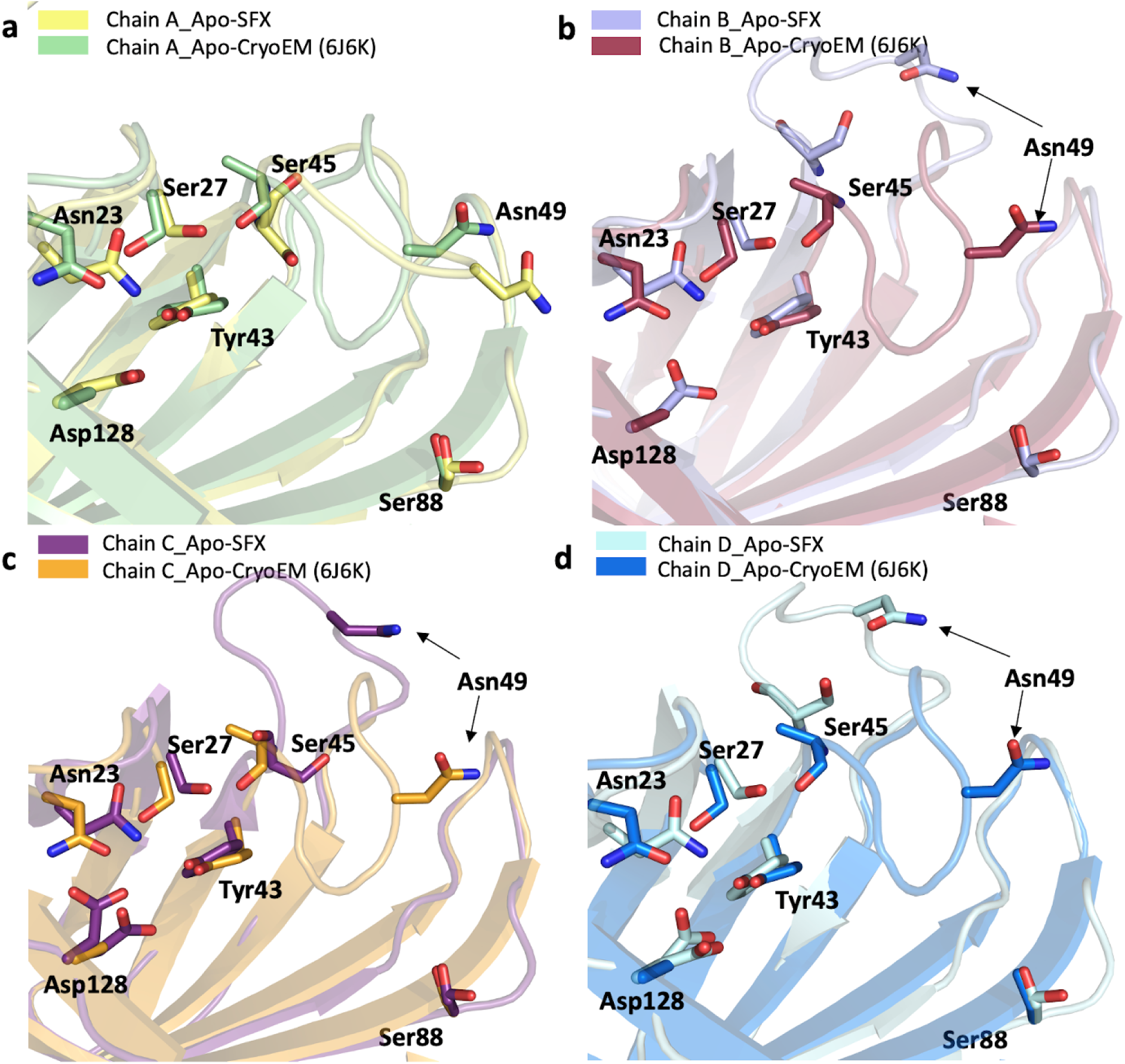
Biotin binding site of Apo-SFX structure compared with Apo-CryoEM (PDB ID: 6J6K). Chain A-D of Apo-SFX is superposed with streptavidin structure (PDB ID:6J6K) in panel a-d, respectively (Supp Table 2) Conformational changes are observed on the L3/4 region where the residue 49 is indicated with black arrow for each chai compared to apo-state streptavidin (PDB ID:6J6K).

**Supplementary Fig. 9:**
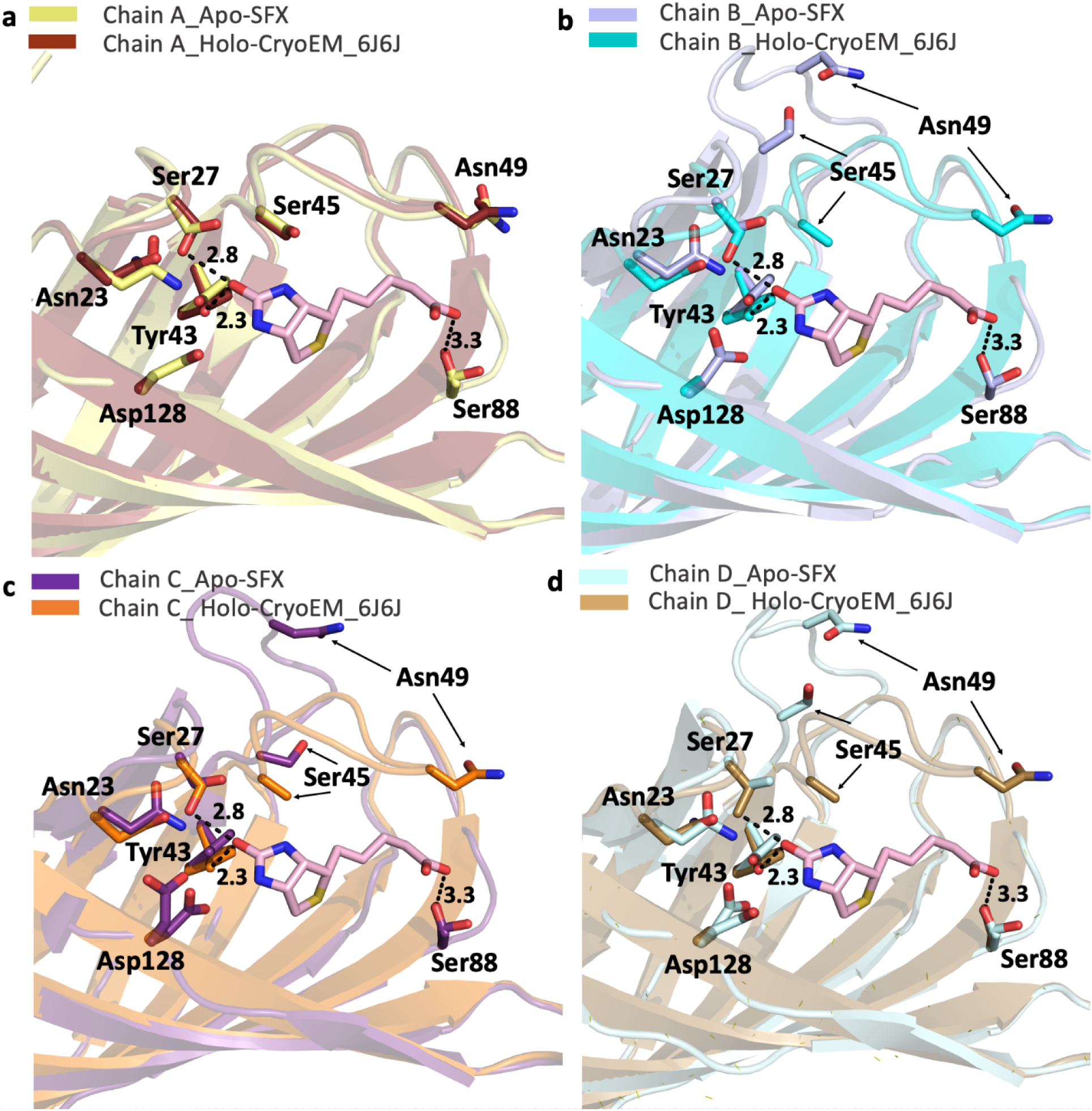
Superposition of our Apo-SFX structure and Holo-CryoEM structure of streptavidin in complex with biotin (PDB ID: 6J6J). a) Superposition of Chain A of Apo-SFX and biotin-bound (PDB ID: 6J6J) streptavidin structures with a RMSD of 0.38 Å. **b)** Superposition of Chain B of two streptavidin structures has an RMSD of 0. 41 Å. **c)** Superposition of Chain C of two structures with a RMSD of 0. 41 Å. **d)** Superposition of Chain D of two structures with a RMSD of 0. 37 Å.

**Supplementary Fig. 10:**
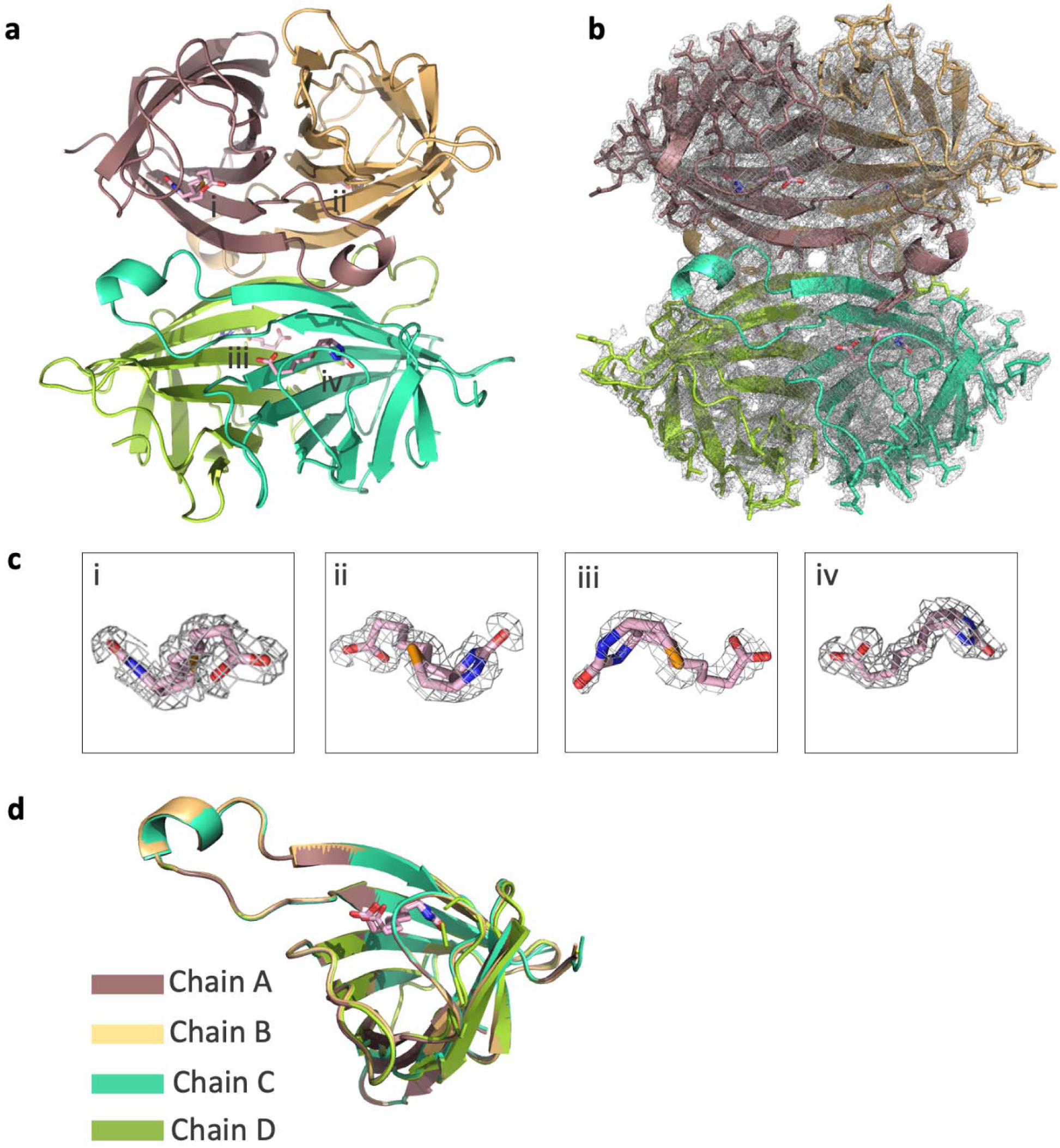
Holo-SFX structure bound with selenobiotin (PDB ID: 5JD2). **(a)** Holo-SFX structure is colored based on chain. **(b)** 2*F*o-*F*c simulated annealing-omit map at 1 sigma level is colored in gray. **(c)** 2*F*o-*F*c simulated annealing-omit map of four selonobiotins (light pink) at 1 sigma level are colored in gray. **(d)** Each chain of streptavidin is superposed with an overall RMSD of 0.13 Å.

**Supplementary Fig. 11:**
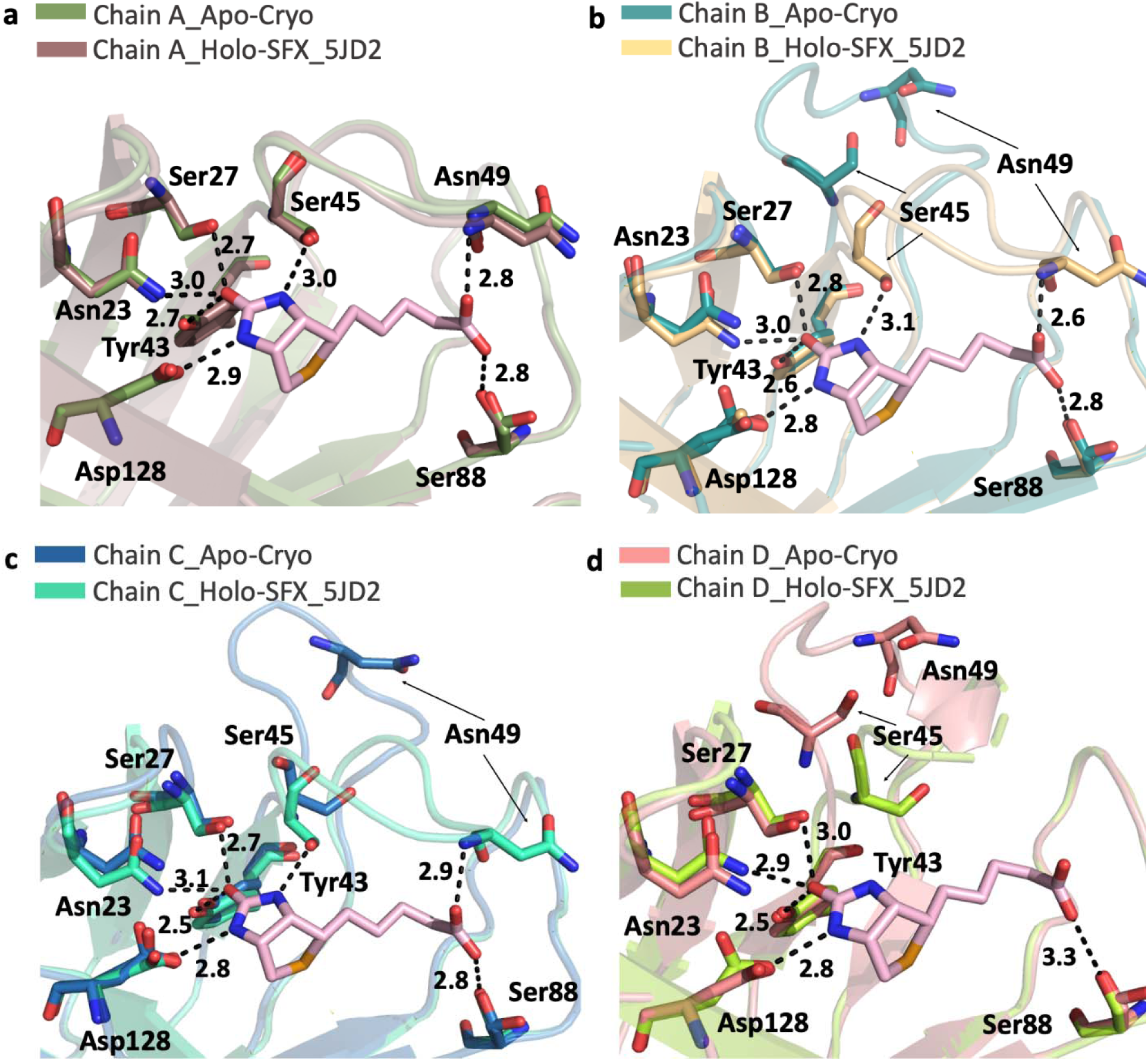
Superposition of Apo-Cryo structure and Holo-SFX structure (PDB ID: 5JD2) around the binding pocket. **(a)** Chain A of both streptavidin structures is superposed with a RMSD of 0. 14 Å. **(b)** Chain B of both streptavidin structures is superposed with a RMSD of 0.14 Å. **(c)** Chain C of both streptavidin structures is superposed with a RMSD of 0.24 Å. **(d)** Chain D of both streptavidin structures is superposed with a RMSD of 0.19 Å. Selenobiotin is colored in light pink and hydrogen bonds are shown with dashed lines.

**Supplementary Fig. 12:**
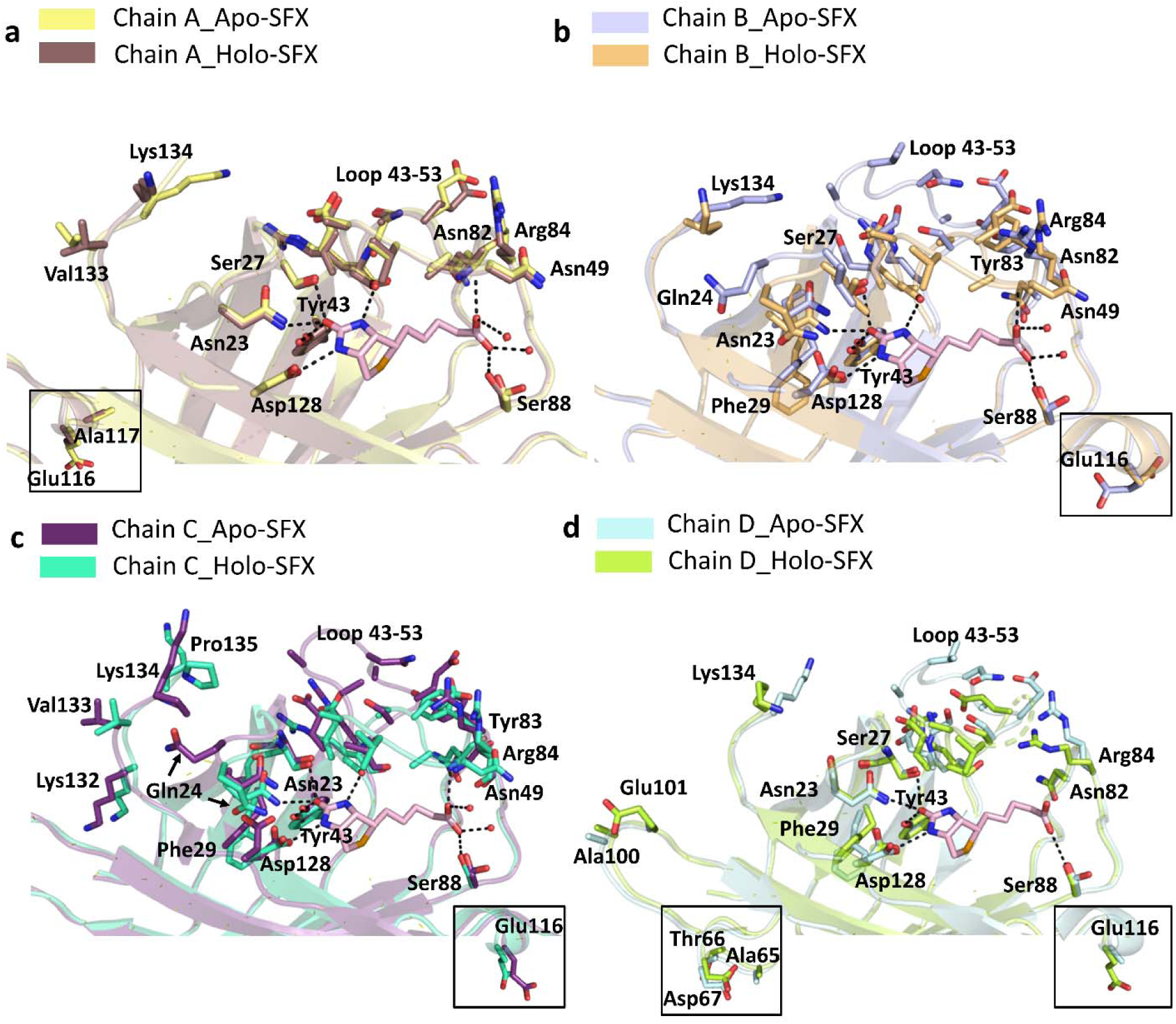
Side chain conformational differences, which were obtained by overall superposition of Apo-SFX and Holo-SFX (PDB ID:5JD2) structures of streptavidin, display effect of ligand binding. Sidechains with different conformations were shown with sticks and labeled in panel a-d. Residues which are apart from the binding side were represented in the boxes. Each chain was colored according to the previous figure legends. Ligand interactions were shown by black dashed lines.

**Supplementary Fig. 13:**
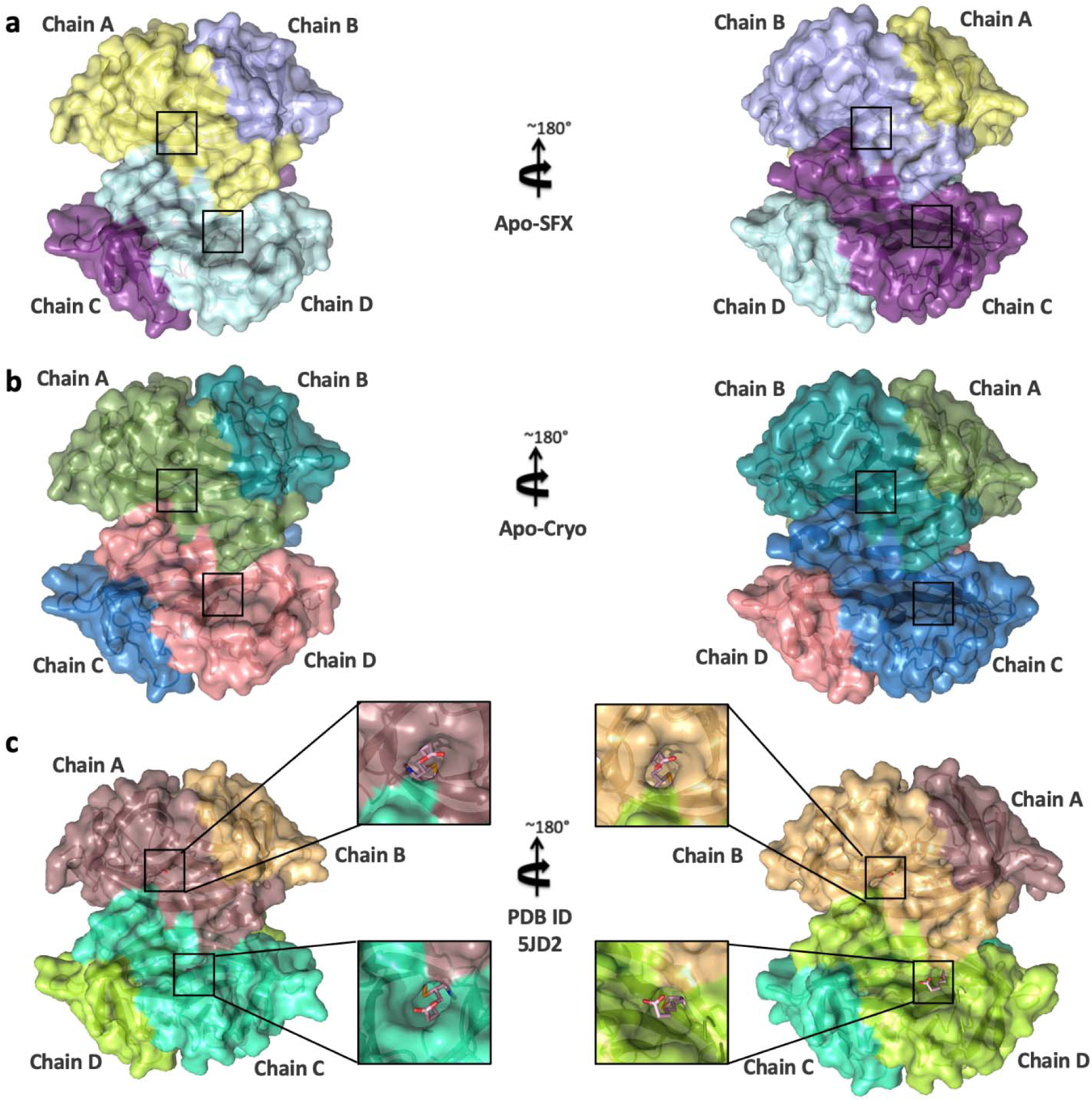
Surface representation of streptavidin structures. **(a)** Apo-SFX structure of streptavidin **(b)** Apo-Cryo structure of streptavidin **(c)** Holo-SFX structure (PDB ID: 5JD2) are colored based on chain. “The binding pocket for selenobiotin which is colored in light pink is indicated with black squares in the panels.”

**Supplementary Fig. 14:**
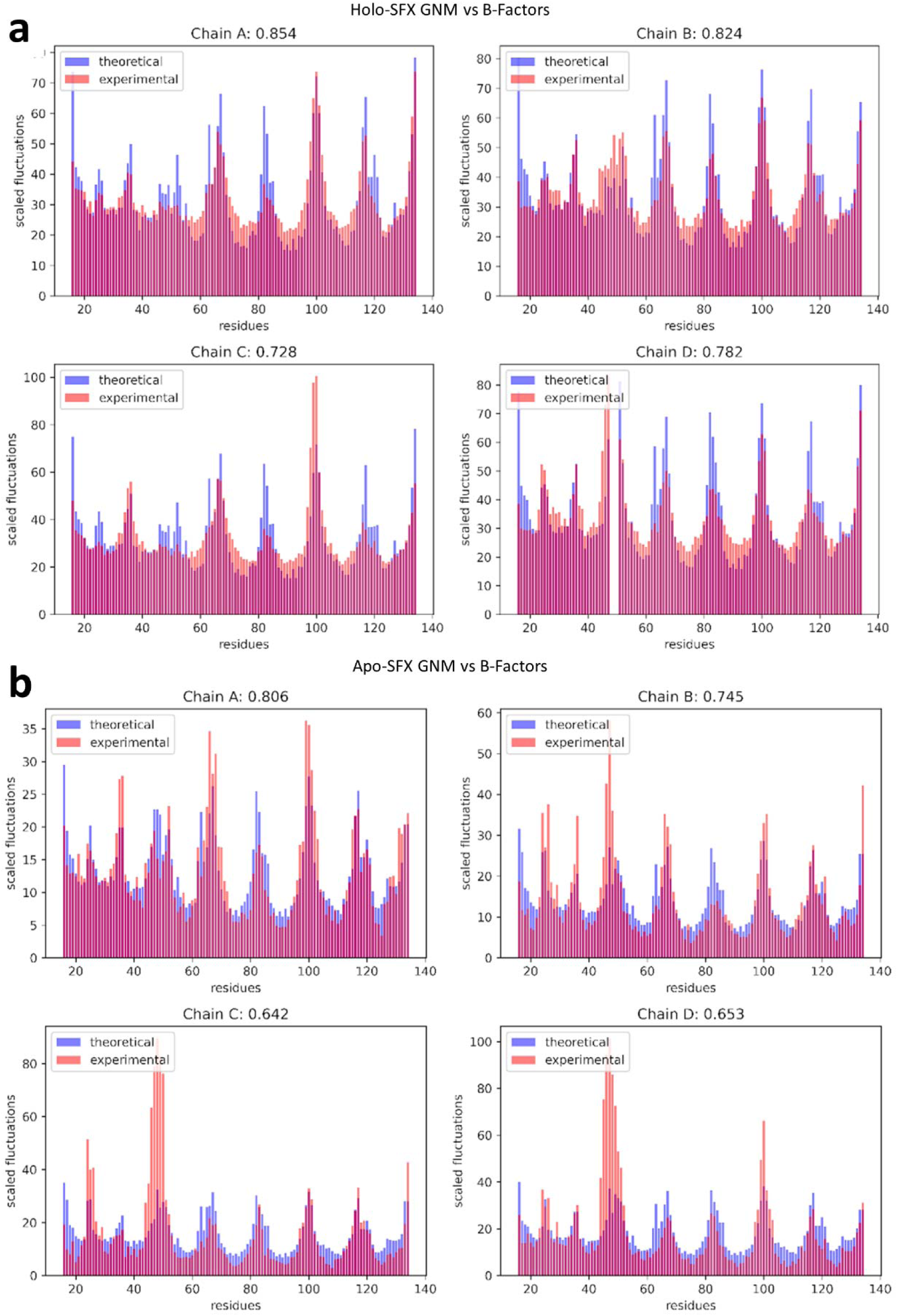
Theoretical residue fluctuations from GNM compared with the experimental fluctuations. The correlations between the calculated GNM fluctuations (theoretical) and the B-factors (experimental) are provided for each chain respectively in the results of both Holo-SFX (a) and Apo-SFX (b) structures.

**Supplementary Fig. 15:**
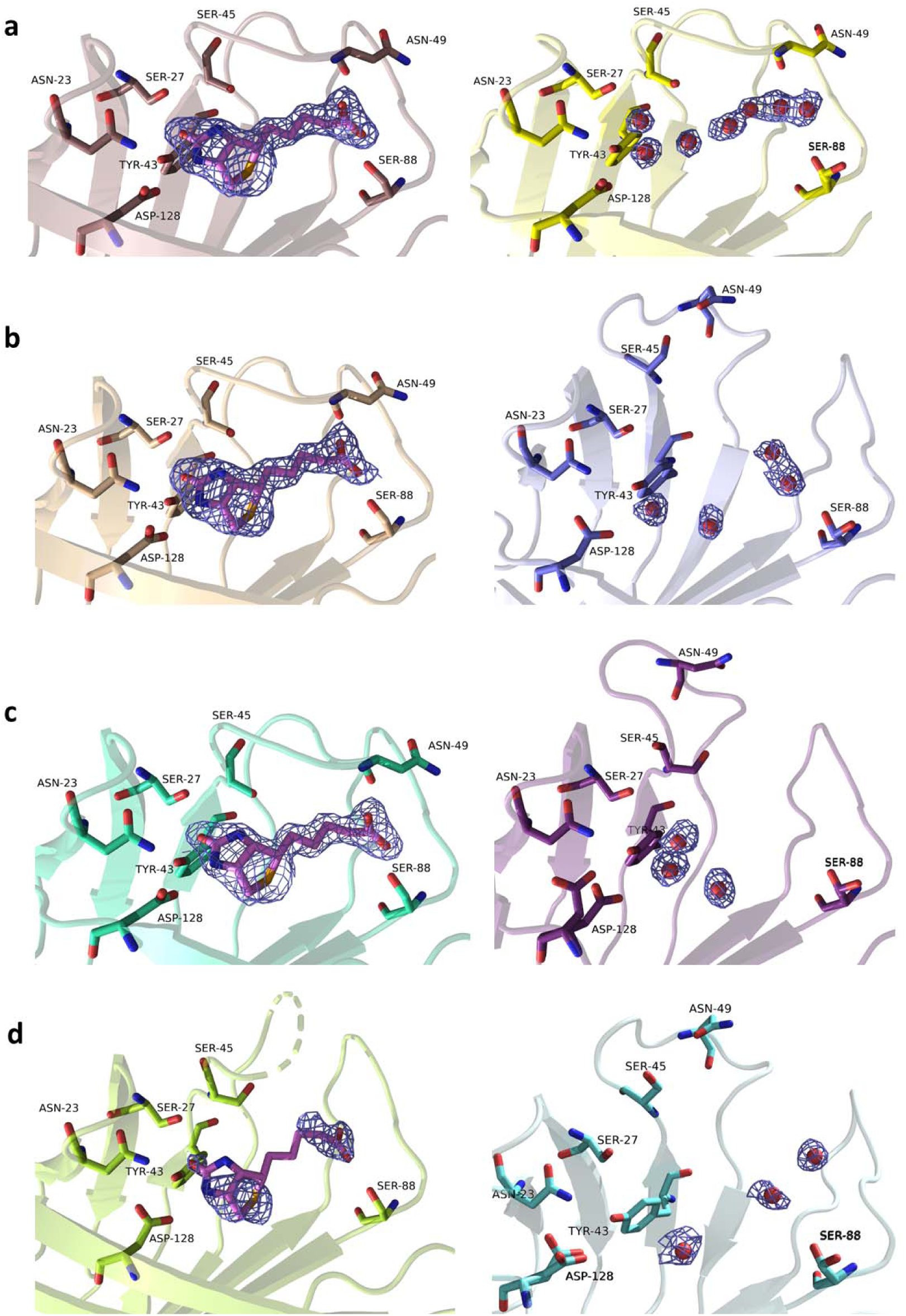
Electron density maps of all ligands of Holo-SFX (5JD2) and corresponding water molecules at the binding site of Apo-SFX structures. Electron densities derived from a 2*F*o-*F*c at 1 sigma level and colored in slate. The electron density map of water molecules in the A chain is similar to the selenobiotin ligand electron density map with continuous electron clouds. From the A chain to the D chain (panel a-d), the water-binding activity of the binding pocket of each subunit is asymmetrically different compared to selenobiotin-bound streptavidin corresponding subunits.

**Supplementary Fig. 16:**
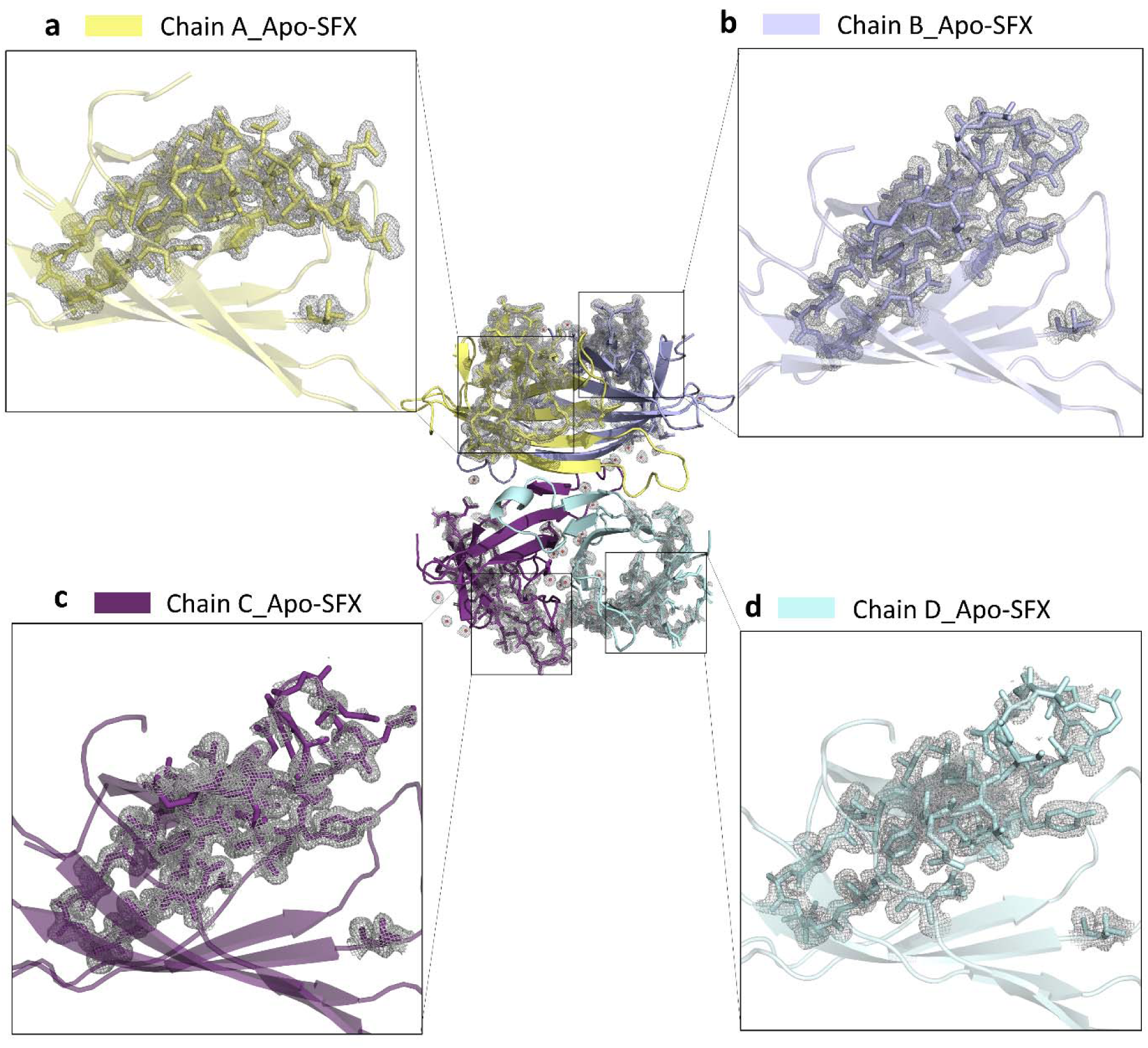
Electron density map of residues which is located near the binding site of the Apo-SFX structure of streptavidin. 2*F*o-*F*c simulated electron density map is colored in slate in panel a-d. Chain colors are presented as described before. Chain A and B have great overlap with electron density maps. However, some amino acid residues which are on loop with binding site residues are not corresponded with electron density maps in Chain C and Chain D which are disoriented in previous structures.

**Supplementary Fig. 17:**
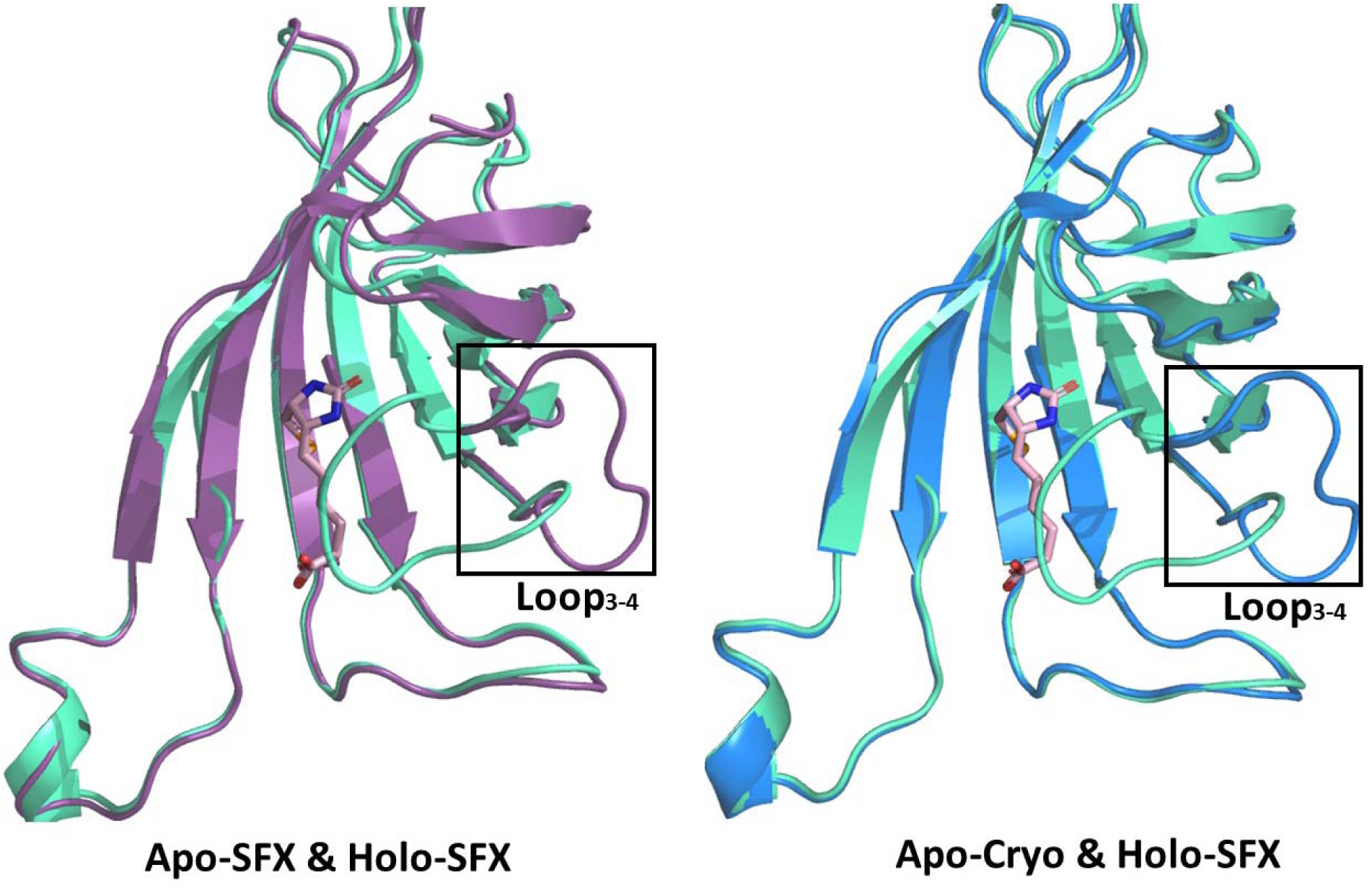
Representation of loop-closed and loop-open “lid” conformations in superimposed C-chains of the streptavidin. On the left, loop-open conformations are colored in violet-purple and marine for Apo-SFX in left and Apo-Cryo structures in right, respectively. On the right, loop-closed conformation is colored in green-cyan in Holo-SFX for each C-chains of Apo-SFX structure.

**Supplementary Table 1:** Root mean square deviation (RMSD) values between streptavidin structures. RMSD values for the L3/4 (residues 45-52) are indicated in the parentheses.

| RMSD (Å) of Cα<sup>†</sup> |  |  |  |  |
| --- | --- | --- | --- | --- |
|  | Chain A<br>Apo-SFX | Chain B<br>Apo-SFX | Chain C<br>Apo-SFX | Chain D<br>Apo-SFX |
| Chain A<br>Apo-Cryo | 0.214<br>(0.135) | - | - | - |
| Chain B<br>Apo-Cryo | - | 0.219<br>(0.119) | - | - |
| Chain B<br>Apo-Cryo | - | - | 0.228<br>(0.148) | - |
| Chain B<br>Apo-Cryo | - | - | - | 0.220 (0.478) |
| Chain A<br>6J6K | 0.426<br>(2.879) | - | - | - |
| Chain B<br>6J6K | - | 0.457<br>(4.484) | - | - |
| Chain C<br>6J6K | - | - | 0.481<br>(3.276) | - |
| Chain D<br>6J6K | - | - | - | 0.432 (3.649) |
| Chain A<br>6J6J | 0.389<br>(0.341) | - | - | - |
| Chain B<br>6J6J | - | 0.414<br>(5.076) | - | - |
| Chain C<br>6J6J | - | - | 0.411<br>(2.975) | - |
| Chain D<br>6J6J | - | - | - | 0.375 (5.060) |
| Chain A<br>5JD2 | 0.269<br>(0.165) | - | - | - |
| Chain B<br>5JD2 | - | 0.213<br>(4.839) | - | - |
| Chain C<br>5JD2 | - | - | 0.360<br>(2.889) | - |
| Chain D<br>5JD2 | - | - | - | 0.262 (2.112) |

**Supplementary Table 2:**
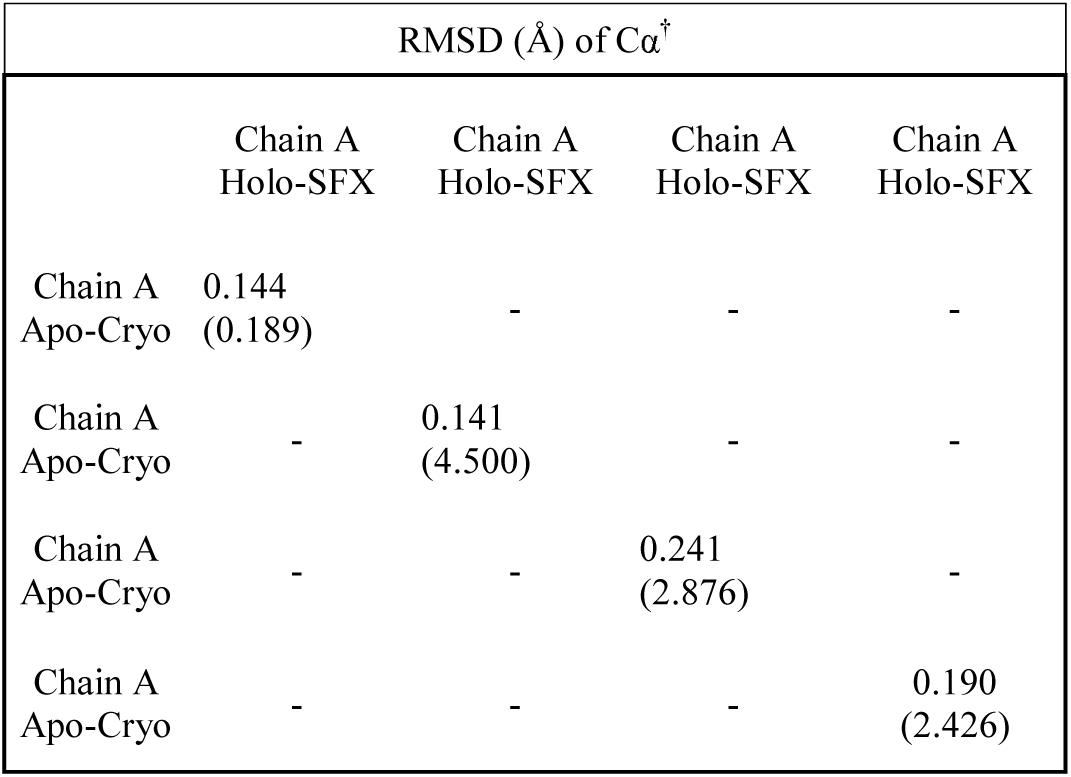
Root mean square deviation (RMSD) values between Apo-Cryo and Holo-SFX streptavidin (PDB ID: 5JD2). RMSD values for the L3/4 (residues 45-52) are indicated in the parentheses.

